# Missing data in single-cell transcriptomes reveals transcriptional shifts

**DOI:** 10.1101/2025.08.15.669765

**Authors:** Ruizhe Chen, Yu-Che Chung, Beth Kelly, Hannah Moore, Sanjib Basu, Dmitrijs Lvovs, Paul M. Gueguen, David E. Sanin

## Abstract

Profiling thousands of single cell transcriptomes is routine, yet cell prioritization based on response to biological perturbations is challenging and confounded by clustering, normalization and dimensionality reduction strategies. We developed a scoring approach independent of these obstacles that unbiasedly identifies distinct transcriptomes within a set based on missing data patterns, allowing cell prioritization and feature selection for downstream analysis. Our method applied to *D. discoideum* reveals a metabolic shift that marks the transition between the amoeboid and aggregated states of this model organism.

## Main

Single cell transcriptomes are dominated by zero-counts. This may be attributable to technology specific artifacts^1–3^, but recent analyses suggest that this phenomenon results primarily from low-expression and limited transcript capture^4^. Traditionally, zero-counts are seen as a barrier that must be resolved computationally, say via imputation^5^, or with technological developments. However, zero-counts can define highly variable features^6^ and identify cell types^7^. Our study builds on this small but growing body of work by adapting statistical methods commonly used to evaluate patterns of missing data in low-dimensional clinical datasets to single cell RNA sequencing (scRNAseq) settings. Specifically, we apply the concepts of data ‘missing at random’ and ‘observed at random’, which distinguish ordinary, random lapses in data collection from more interesting and potentially problematic scenarios that arise when missingness is confounded with variables of interest. For instance, one may detect survey biases introduced by participants’ behaviours that result in missing data^8^. Patterns of missingness in high-dimensional scRNA data are admittedly different in both cause and consequence from those seen in low-dimensional clinical studies. Specifically, we expect that occurrences of zeros in single cell data are not random, but rather vary systematically across cell types and experimental conditions. Nor are zero-counts strictly ‘missing’, as it is not possible to distinguish true absence from not recorded. Nonetheless, it has been our experience that evaluating the extent to which scRNAseq data appears to be ‘Observed At Random’ (OAR) using zero-counts to define missing data patterns (MDPs) provides a unique and powerful tool for characterizing transcriptional patterns, and identifying shifts and deviations therefrom.

In the following paragraphs we describe our methodology (OAR scores) to perform cell prioritization and feature selection for downstream analysis. We discuss treating zero-counts as missing data and describe the statistical framework that allows us to examine these patterns in detail, illustrating the implementation and interpretation of our method across 21 datasets. We later focus our analysis on *Dictyostelium discoideum* to highlight novel biology that can be discovered with our method. Typically, cell prioritization and differential gene expression analysis relies on cluster labels^9–11^, dimensionality reduction techniques^12^ and normalization strategies^13^, which can confound the effect of biological perturbations. Our method is independent of all these factors, providing an avenue to extract meaningful biological insights from the thousands of single cell transcriptomes that are routinely generated.

The core of our method examines the extent to which a cell approaches the OAR assumption^14^, that is, the extent to which the MDPs fail to explain observed gene expression in each cell. OAR scores are calculated in two broad steps (**Fig. 1a**): First, MDPs are defined by identifying co-expressing gene modules based on a binary classification of expressed (i.e. greater than 0 counts or ‘observed’) or not-expressed (i.e. 0 count or ‘missing’) genes in each cell (**Fig. 1a**). MDPs group genes that co-occur in cells, setting a conservative tolerance to allow for non-exact matches. Second, transcriptional shifts are scored in individual cells where the MDPs are less effective at predicting ‘observed’ values (**Fig. 1a**). We propose that MDPs (i.e. conserved instances of zero-counts across cells), are essentially co-expression patterns related to cellular identity and function. Consequently, when MDPs that are defined based on a population of cells insufficiently explain the observed values in a given cell, we infer a fundamental shift in the transcriptional program of that cell and highlight it by assigning it a high OAR score (**Fig. 1a**). Thus a cell with a high OAR score (i.e. approaching the OAR assumption) is distinct from all the others used to define the MDPs in the first place. In this way, our method allows for the rapid identification of transcriptional shifts arising from biological perturbations independently of differential gene expression testing.

**Fig. 1.**
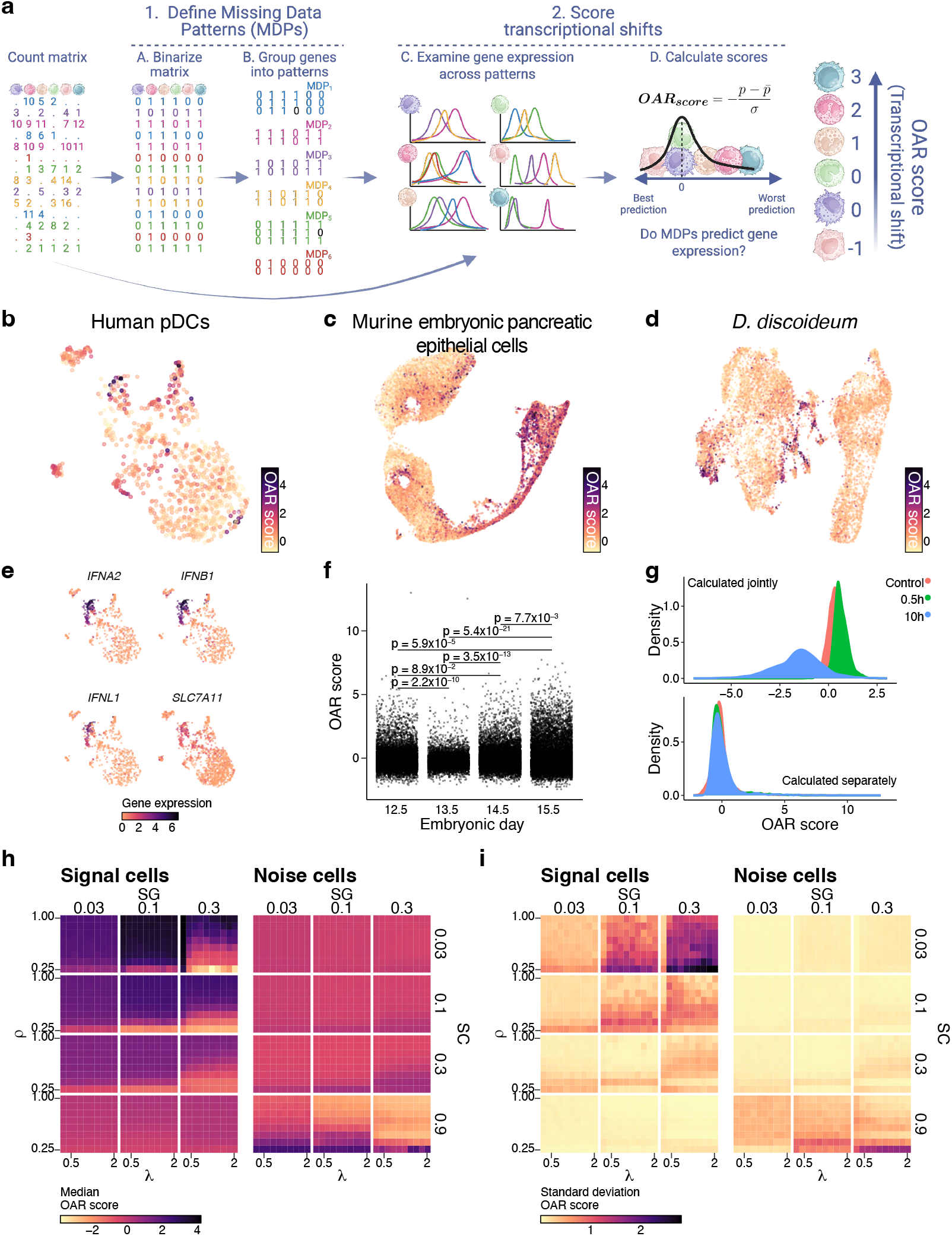
**a**, Schematic overview of OAR score calculation. **b-g**, OAR scores from naive and activated human plasmacytoid dendritic cells (pDCs, **b**,**e**; GSE157305), developing murine embryonic pancreatic epithelial cells (**c**,**f**; GSE132188), and vegetative plus starved *Dictyostelium discoideum* cells (**d**,**g**; GSE164010), visualized as a Uniform manifold approximation and projection (UMAP) colored according to OAR scores (**b-d**). **e**, Expression of indicated genes in pDCs shown as a UMAP. **f**, OAR scores compared across embryonic days in pancreatic epithelial cells. Adjusted p-values contrasting indicated groups are shown from a Kruskal-Wallis test followed by a Dunn’s test with a Benjamini-Hochberg correction. **g**, OAR scores calculated on all cells simultaneously (top) or separately by biological condition (bottom) are shown. **h**,**i**, Median OAR score (**h**) and standard deviation (**i**) for simulated signal and noise cells across indicated values for gene expression correlation (ρ), rate (λ), fraction of signal genes (SG), and fraction of signal cells (SC). Each simulation consisted of 500 cells, with 5000 genes. Median and standard deviation were calculated across 100 replicates per combination of indicated parameters.

We illustrate the results obtained from applying this method using several publicly available datasets (**Fig. 1b-g**, **Extended Data Fig. 1, Supplementary Fig. 1-4**). For instance, in a dataset of human plasmacytoid dendritic cells (pDCs), high OAR scores coincided with highly activated pDCs expressing effector molecules typical for these cells^15^ (**Fig. 1b,e**). Likewise, high OAR scoring cells in murine embryonic pancreatic epithelial cells were associated with embryonic development (**Fig. 1c,f**). Notably, we observed that in *D. discoideum* cells transitioning from the amoeboid to the aggregated state, OAR scores were affected by the biological condition in the underlying data (**Fig. 1g** - top panel and **Supplementary Fig. 3d**,**e**). We hypothesised that the identified MDPs (**Supplementary Fig. 3b**) failed to capture meaningful co-expression patterns in this setting when defined across all the cells simultaneously. We therefore calculated OAR scores for each biological condition separately (**Fig. 1g** - bottom panel and **Supplementary Fig. 4**), resulting in OAR scores comparable across conditions (**Fig. 1d,g**). In this manner, our methodology can reveal biological effects that produce spurious co-expression patterns in addition to cells with distinct transcriptional programs.

OAR scores were not associated with low abundance gene filtering (**Supplementary Fig. 5**), genes associated with tissue digestion (**Supplementary Fig. 6a**,**b**) or low quality cells produced due to sample processing (**Supplementary Fig. 6-7**). scRNAseq protocols produce different proportions of missing values, and if tested together, those differences are reflected in test results (**Supplementary Fig. 8a**,**b**). However, analogous to what we observed with *D. discoideum* (**Fig. 1g** and **Supplementary Fig. 3-4**) calculating scores for each technology separately, produced comparable OAR scores (**Supplementary Fig. 8c**,**d**), which were robust across batches and donors. Thus, our method can rapidly reveal confounding factors driving diverging MDPs, allowing us to prioritize cells with transcriptional shifts both across and within biological conditions (**Fig. 1b-g**).

We reasoned that if a high OAR score reflects a shift in transcriptional programs, we could reduce high scores by removing the genes associated with these changes. For that purpose we defined ‘signal cells’ as those with an OAR score greater than 2, and subsequently we performed a Wilcoxon Rank Test to identify differentially expressed genes (adjusted p-value < 0.1; log_2_ fold change > 0) in these cells compared to all others and designated these as ‘signal genes’. OAR scores in signal cells calculated on the full gene expression matrix were significantly higher than those calculated on a matrix without signal genes in all datasets examined (**Extended Data Fig. 2**). This supports our view that high OAR scores are associated with transcriptional shifts. To further demonstrate this property of our test, we simulated scRNAseq data by implanting ‘signal genes’ in ‘signal cells’ with a distinct correlation structure employing a Gamma-Poisson distribution (**Extended Data Fig. 3a**), and confirmed that for a wide range of simulation parameters and signal gene and cell proportions, we obtained high OAR scores for signal cells (**Fig. 1h,i**, **Extended Data Fig. 3b-d**). Notably, increasing the proportion of ‘signal cells’ (e.g. to 90% of tested samples) increases OAR scores in ‘noise cells’ (i.e. not ‘signal cells’). This is consistent with the idea that our test highlights cells with a distinct transcriptional program compared to the majority of cells tested, which would be the case for ‘noise cells’ when they are a minority in the expression matrix. Interestingly, in both real and simulated data, high OAR scores are associated with high percentages of missing values (**Extended Data Fig. 3c, Supplementary Fig. 1d, 2d, 4**). We examined this in detail by simulating data with greater sparsity in ‘noise cells’ and comparing resulting OAR scores (**Extended Data Fig. 4**). We found that matching or greater sparsity in ‘noise cells’ compared to ‘signal cells’ decreases OAR scores in the latter, which suggests that our test is best suited to identify cells with a distinct transcriptional program and generally greater sparsity.

**Fig. 2.**
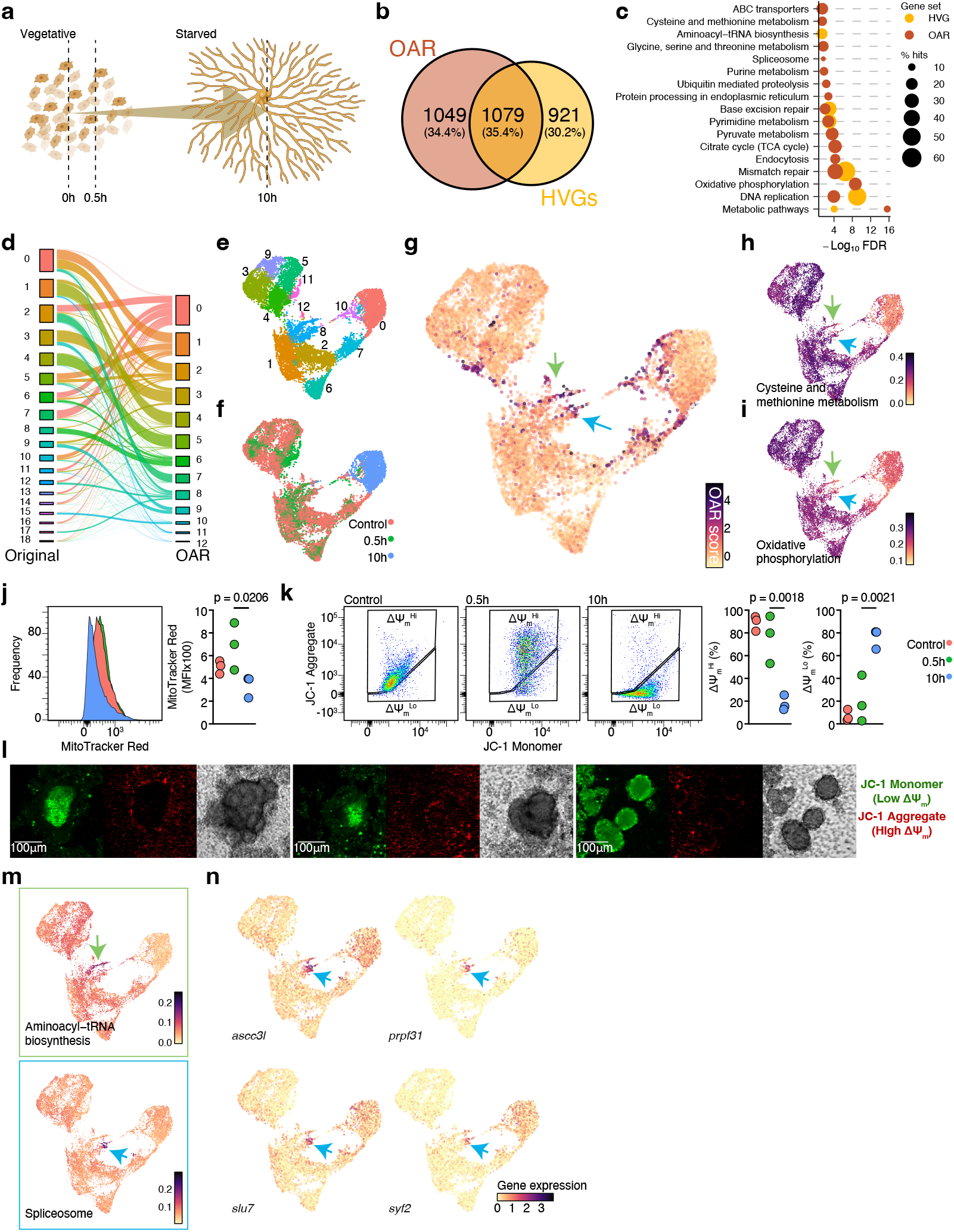
**a**, *D. discoideum* experimental overview with sampling points indicating when scRNAseq was performed. **b**, Overlap between highly variable genes (HVG - gold) and OAR associated genes (brown). **c**, Pathway enrichment analysis against kyoto encyclopedia of genes and genomes (KEGG) pathways for HVG (gold) and OAR associated genes (brown), showing percent of pathway covered (% hits) and enrichment significance. **d**, Alluvial plot relating original (left) and OAR associated gene-derived (right) clustering. **e-i**, UMAP with cells colored by cluster (**e**), sampling point (**f**), OAR scores (**g**) or calculated enriched pathway scores, highlighting regions of interest (green and blue arrows) (**h-i**). **j-l**, *D. discoideum* cells were examined for mitochondrial activity with mitochondrial membrane potential (ΔΨ_m_) sensing probes via flow cytometry (**j**,**k**) or microscopy (**l**). Cells were starved for indicated times (**j**,**k**) or for 24h (**l**) prior to acquisition. Decreased ΔΨ_m_, evident as lower MitoTracker Red fluorescence or increased JC-1 monomers, was observed after 10h of starvation (**j**,**k**). Cells forming multicellular aggregates had consistently depolarized mitochondria, whereas cells in the periphery displayed high ΔΨ_m_ (**l**). Results are representative of 2 independent experiments with individual replicates as points. ANOVA followed by unpaired T-tests were used to define statistically significant differences between experimental groups. **m**,**n**, UMAP with cells colored by calculated enriched pathway scores (**m**) and indicated gene expression (**n**), highlighting cells of interest (green and blue arrows).

Our method produces a quantitative measure of transcriptional drift that is associated with a set of genes expressed in the cells with a high OAR score. We sought to exploit this by identifying genes that were associated with this score using a likelihood-ratio test^16^ with an overly stringent false discovery rate threshold. We found that using natural splines we were able to detect thousands of genes that were significantly associated with OAR scores (**Extended Data Fig. 5**). Moreover, these OAR associated genes only partially overlaped with highly variable genes (HVGs) across three datasets tested (**Fig. 2a,b**, **Extended Data Fig. 5**). Notably, despite being defined in a cluster agnostic manner, OAR associated genes could partly recreate the cell clustering obtained from a typical scRNAseq analysis pipeline (**Supplementary Fig. 9**). To further understand the biological implications of these gene sets, we focused on the *D. discoideum* dataset (**Fig. 2a**) and performed pathway enrichment analysis. We discovered that the transcriptional programs overrepresented in each gene set only partly overlapped (**Fig. 2c, Extended Data Fig. 6**), and found that metabolic pathways were particularly enriched in OAR associated genes, and not HVGs (**Fig. 2c, Extended Data Fig. 6b**), reflecting the biological perturbation studied where cells are nutrient-starved to induce aggregation (**Fig. 2a**). We next used OAR associated genes to re-calculate the PCA, UMAP and clustering of *D. discoideum* cells, and found that although some overlap in the results was evident (**Fig. 2d**), OAR associated genes separated each biological condition more clearly (**Fig. 2e,f**, **Extended Data Fig. 7**), with high OAR scoring cells clustering together (**Fig. 2g**). Notably, we previously reported that cysteine and methionine metabolism (**Fig. 2h**) and oxidative phosphorylation (**Fig. 2i**) were critical for *D. discoideum* aggregation^17^, and our current analysis orthogonally revealed that feature of this process (**Fig. 2h,i**). Moreover, when we mapped the pathways enriched in OAR associated genes to the new UMAP (**Fig. 2g-i**, **Extended Data Fig. 6c**), our results made new predictions which could be validated experimentally. First, our analysis reveals that in nutrient replete *D. discoideum* cells, a transcriptional shift away from oxidative phosphorylation is already evident (**Fig. 2g,i**, green and blue arrows, **Supplementary Fig. 10a**). We hypothesized that these cells could be pioneers in forming cellular aggregates and confirmed this with flow cytometry and live microscopy of *D. discoideum* cells labelled for mitochondrial activity (**Fig. 2j-l**, **Extended Data Fig. 8, Supplementary video 1**,**2**). Moreover, we observed highly specific transcriptional programs occurring in high OAR scoring cells, including Aminoacyl−tRNA biosynthesis (**Fig. 2m**, green arrow, **Supplementary Fig. 10b**), as well as spliceosome activation (**Fig. 2m**, blue arrows, **Supplementary Fig. 10c**) associated with highly expressed splicing machinery (**Fig. 2n**, blue arrows). Thus OAR scores and associated transcriptional shifts revealed new insights into the dynamic differentiation process of *D. discoideum* cells.

OAR score calculations are computationally efficient (**Extended Data Fig. 9**), make minimal assumptions about data structures or protocols, and are independent of clustering, normalization and dimensionality reduction techniques. Close examination of results can reveal confounding variables in the analysis of single cell transcriptomes. Critically, OAR scores can be applied broadly to discover biologically meaningful transcriptional shifts, robustly across batches, technologies and organisms. We have identified 2 limitations to OAR scores. First, they are influenced by the cells tested, as MDPs are by necessity defined across sets of cells, and consequently OAR scores will vary depending on the samples included in the analysis. However, these results will be contextually meaningful. Second, OAR scores are somewhat restricted to transcriptional shifts occurring in cells with a higher degree of sparsity. This phenomenon could be similar to the observation that cell differentiation often results in a restricted transcriptional diversity^18^. Future studies will focus on understanding this relationship. Currently, OAR scores offer analysts the possibility of focusing their study on a divergent population of cells that may be extracted from a set of transcriptomes without specialized knowledge or cell labels, potentially streamlining analysis and subsequent real-world validation.

## Acknowledgements

We would like to thank Drs. Leslie Cope, Elana Fertig, Ravi Varadhan, Katarzyna Grzes, Edward Pearce and Erika Pearce for their insightful comments and suggestions as we developed this methodology. We would also like to thank all the members of the Pearce laboratories past and present for generating some of the key datasets contained in this work. D.E.S. is supported by the Prostate Cancer Foundation (22YOUN16) and the National Institutes of Health (R21CA273501).

## Methods

### Zeros as missingness in scRNAseq data

Single cell transcriptomes are dominated by zero-counts. Although these values are often referred to as ‘dropouts’, they are expected given that count data follows a Gamma-Poisson distribution^4^, reflecting the biology of the cells studied and the overall expression of a gene at any given time. Several studies have sought to make a distinction between ‘technical zeros’, genes with no expression in several cells and relatively high in other cells, and ‘biological zeros’, which are zero because the gene is not expressed. However, dropouts, which led to the implementation of zero-inflated distributions to model single cell transcriptomes^1^ and concerns of technical cell-to-cell variability confounding biological insights^2^, are mainly limited to technologies that do not use unique molecular identifiers^3^. Thus, zero-counts in single cell data may be studied to understand the biological properties of cells. This aspect of scRNAseq data has been used in the past to define highly variable features^6^, and identify cell types^7^. Here we use zero-counts to find co-expression patterns and subsequently define cells with divergent transcriptional programs that warrant further characterization.

Missing data, which is to say when a variable is not observed due to an experimental limitation, is a frequent problem that statistics must address. In scRNAseq applications, zero-counts arise from a relationship between the expression level of a gene and the technical limitations in capturing all expressed transcripts. Thus, a zero-count in scRNAseq data may be understood as missing as it is not possible to distinguish what is truly no expression versus no capture.

### The Observed at Random Assumption

We previously developed a diagnostic test to assess the underlying missing data mechanisms in incompletely observed patient-level clinical data^19^, which is essential for applying proper statistical inference methods. A missing data mechanism is a probability model that specifies the relationship between the values and missingness of a random variable by treating missing data indicators themselves as random variables^14,20,21^. Here, we study one such mechanism in scRNAseq data, specifically the OAR assumption^14^, under which the conditional probability of missingness functionally does not depend on the observed values^22^. In order to describe the ability of our methodology to test for OAR in scRNAseq data, it is necessary to define other missing data mechanisms that could be at play and impact the interpretation of our results. These are:

i. When the missingness does not depend on the unobserved values, the missing data is said to be missing at random (MAR).
ii. When the missingness depends on the unobserved values, the missing data is said to be missing not at random (MNAR).
iii. When the missing data is MAR and observed data is OAR, the missingness is completely at random (MCAR)^23^. That is, the probability of observing the missing data indicators given the data, is equivalent to the probability of observing the missing data indicators without conditioning on the data.

We previously demonstrated that our testing framework for MCAR in mixed-type datasets can effectively examine the OAR assumption^19^. Intuitively, the MAR component of MCAR can not be tested without imposing external restrictions^24,25^. This is because it is difficult to establish the association of missingness with values that are not observed. In fact, it is the OAR assumption component of MCAR that is testable, and thus it is the core of our missing data diagnostic test. From the OAR assumption it follows that the distribution of the observed components do not depend on the MDPs^19^, which is what our method examines. However, if the observed values and unobserved values are stochastically dependent, then they are “connected”. Since the MDPs can functionally depend on the unobserved values under OAR, this dependence creates a connection from observed values to MDPs through unobserved values. Consequently, provided not all columns (cells) are independent, which can be reasonably assumed in scRNAseq data, our test can also detect MNAR. Thus, our diagnostic test has two detection regions^19^ (i.e. two sets of conditions where the null hypothesis is rejected and the p-values are small): 1) Observed values are not OAR; 2) Missing values are MNAR, with observed values correlated.

### OAR test for High-Dimensional Count Data

OAR scores are a result of testing the homogeneity assumption among the observed data grouped by MDPs^19^. A distinctive feature of our test lies in that it associates the missingness patterns with observed values instead of relying on missingness patterns alone. In the case of scRNAseq, there is a pressing need to understand what missing data implies about the biology of the cells studied. We have thus adapted our previous findings to use the resulting test statistics as distinctive cell signals. This is achievable because our test works as an iterative procedure. Instead of assuming a multivariate normal distribution and yielding a single statistic assessing the overall level of departure from the null^26^, our test yields single statistics (p-values) for each cell (column) that allow for individual interpretations. Given the OAR assumption holds, the observed distributions of the observed values should be identical within groups defined by distinct MDPs. Notably, the null OAR assumption does not hold in all examined scRNAseq datasets. However, cells still exhibit different levels of departure from the null, with some cells approaching closer to the null (OAR) and thus revealing transcriptional alterations.

Previously, we defined MDPs in mixed-type clinical data using exact matching^19^. However, scRNAseq data comprises tens of thousands of features across tens (or hundreds) of thousands of cells, yielding few if any exact matches. Therefore, we mitigate the issue of highly sparse MDPs with non-exact matching based on Hamming distance within gene expression deciles (**Supplementary Fig. 1-3**). We accomplish this following these steps:

1. We extract the un-normalized count data matrix and remove low abundance genes (i.e. genes expressed in a small fraction of cells). This step is routine in most bioinformatic pipelines, and we found that it had little impact on our results (**Supplementary Fig. 5**). The count matrix is subsequently used to create an indicator matrix where values are either missing (0) or observed (1).
2. The indicator matrix is split into groups of genes expressed in a similar fraction of cells, which achieves two purposes. First, it greatly increases the computational efficiency of the approach as it limits the number of pairwise Hamming distances calculated. Second, genes expressed in a fraction of cells may only be in an MDP with other genes expressed in a similar fraction of cells (i.e. small Hamming distances are only possible between genes expressed by a similar fraction of cells). We chose to split the indicator matrix into 10 bins, corresponding to the 10 quantiles (deciles) calculated across the fraction of cells expressing each gene (**Supplementary Fig. 1-3**).
3. We calculate the Hamming distance between all pairs of genes within each bin using the indicator matrix. Hamming distances are ideally suited to handle binary data and provide an accurate estimator of co-expression for genes across cells. Let the Hamming distance *h* between two genes *a* and *b* expressed across *n* cells be defined as

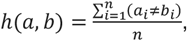

so that *h* = 0 indicates an identical co-expression pattern for genes *a* and *b*, while *h* = 1 represents no co-expression.
4. Let *H*_*i*_ be the pairwise triangular matrix of Hamming distances in bin *i* of the indicator matrix. The values of *H*_*i*_ are normally distributed (**Supplementary Fig. 1a, 2a, 3a**), and we may calculate a tolerance *τ*_*i*_ as

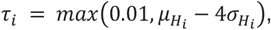

based on the mean *μ* and standard deviation *σ* of *H*_*i*_. Thus *τ*_*i*_ is a value greater than 0.01 for each bin of the indicator matrix.
5. Let *A*_*i*_ be the adjacency matrix for genes in bin *i*, obtained as

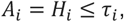

so that a graph *g*_*i*_ may be built consisting of communities of genes (nodes) in bin *i* connected only if their corresponding Hamming distance is below *τ*_*i*_. To maintain only densely connected communities, we examine the maximum eccentricity of *g*_*i*_. If this value is greater than 3, then we adjust *τ*_*i*_ as

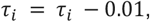

then recalculate *A*_*i*_ and *g*_*i*_ until the maximum eccentricity condition is met. Communities in *g*_*i*_ corresponding to densely connected genes within each bin are defined as MDPs.
6. An MDP vector, indicating what pattern each gene in the count matrix belongs to, is generated by collecting the community identifiers across all bins, with no overlapping identifiers between bins. This approach yields several MDPs (**Supplementary Fig. 1b, 2b, 3b, 4**), yet most genes are not grouped into co-expression patterns (**Supplementary Fig. 5** - bottom panel).
7. We calculate a Kruskal-Wallis (KW) statistic and the corresponding p-value for each cell using all observed gene expression values within that cell with the observations grouped by the MDP vector. We found that treating genes that were not grouped in a co-expression pattern as unique patterns leads to inflated p-values across all datasets examined following the KW test (**Supplementary Fig. 1c, 2c, 3c**). We addressed this p-value inflation by treating these genes as if they all belonged to a distinct co-expression pattern (i.e. no co-expression), in effect reducing the number of tested patterns in each cell. Implementing this approach resulted in low p-values (**Supplementary Fig. 1d, 2d, 3d**) that could be scaled to produce OAR scores.
8. We adjust the calculated p-values for multiplicity using the Benjamini-Hochberg correction, transform the adjusted p-values with a negative natural log and scale this result (subtraction by mean and division by standard deviation). The negative adjusted and standardized p-values are reported as OAR scores.

The p-value yielded from the OAR test on each cell indicates the degree of departure from the null OAR assumption. Larger p-values, with correspondingly higher OAR scores, indicate greater adherence to the OAR assumption by a given cell within the dataset studied.

### Simulation Studies

We applied a Gamma-Poisson hierarchical model in simulating scRNAseq count data. The initial parameters of the considered data generation models were calibrated based on the plasmacytoid dendritic cell (pDCs) dataset, then modified to examine how gene expression distributions impacted the observed results. We divided this data into three submatrices (**Extended Data Fig. 3a**):

1. A submatrix of ‘signal cells’, those that exhibit an OAR score greater than 2, expressing ‘signal genes’, those that can be associated with the resulting scores in ‘signal cells’.
2. A submatrix of ‘noise cells’, those that exhibit an OAR score lower than 2, expressing ‘signal genes’.
3. A submatrix of all cells expressing ‘noise genes’, which are all genes that are not associated with the high OAR scores in ‘signal cells’.

As base simulation parameters we estimated the Pearson correlation matrix and mean gene expression levels using ‘splatEstimate’ from splatter v. 1.28.0^27^ for each submatrix. Then we examined how different correlation structures, cell composition and distribution parameters would affect calculated scores. To accomplish this, Poisson rates were drawn from the underlying Gamma distribution within each submatrix. Then, for each gene, the count data were drawn from a multivariate Poisson distribution^28^ given the assumed between-cell correlation structure. By applying this approach, the missing data is MNAR and the observed data is not-OAR since the missingness probability depends on unobserved and observed data. Iterations, simulated number of cells and number of genes, as well as simulation parameters for correlation, shape and rate are indicated in each figure. Summary statistics and median results across iterations (**Fig. 1h,i**, **Extended Data Fig. 3b**,**c**) or individual iteration results (**Extended Data Fig. 3d**) are shown.

### Bioinformatic analysis

#### Analysis of publicly available single cell RNA-sequencing datasets

Datasets available from public repositories were retrieved (**Supplementary Table S1**), obtaining matrices with raw unfiltered read counts for detected genes in barcoded cells. Read count matrices were processed, analyzed and visualized in R v. 4.4.0^29^ using Seurat v.5^30^ with default parameters in all functions, unless specified. Poor quality cells, with low total unique molecular identifier (UMI) counts and high percent mitochondrial gene expression, were excluded. Filtered samples were normalized and scaled using ‘NormalizeData’ and ‘ScaleData’ to identify highly variable genes using ‘FindVariableFeatures’, blacklisting genes associated with tissue processing based on annotation available in SignatuR v. 0.2.1^31^. Concomitantly, samples were regularized using negative binomial regression (SCTransform)^32^, batch corrected with Harmonyv. 1.2.0^33^ using ‘RunHarmony’ with 30 principal components calculated on the SCT assay using ‘RunPCA’.

Batch corrected dimensionality reductions were used to identify neighbours using ‘FindNeighbors’, cluster cells using ‘FindClusters’ and calculate a Uniform manifold approximation and projection (UMAP)^34^ using ‘RunUMAP’. Resolution for cell clustering was determined by evaluating hierarchical clustering trees and silhouette scores at a range of resolutions (0 - 1.2, with 0.05 step increments) with Clustree v. 0.5.1^35^ and the ‘silhouette’ function in cluster v. 2.1.6^36^, selecting a value inducing minimal cluster instability and a high proportion of silhouette widths above 0. Datasets in Supplementary Fig. 12 were subsetted to include only macrophages, based on the expression of key macrophage markers (*Adgre1, Csf1r, H2*-*Ab1, Cd68, Lyz2, Itgam, Mertk*), retaining only 500 randomly selected cells per biological condition.

#### Identification of signal genes in signal cells

Cells in naive and activated human plasmacytoid dendritic cells (GSE157305), developing murine embryonic pancreatic epithelial cells (GSE132188), and vegetative plus starved *Dictyostelium discoideum* cells (GSE164010) were split into ‘signal cells’, defined as those with an OAR score greater than 2, and ‘noise cells’ for all others. Genes associated with signal cells (signal genes) were then identified by using a Wilcoxon Rank Sum test with ‘FindMarkers’ from Seurat v.5^32^, grouping cells by OAR score status and keeping genes with a positive fold change and an adjusted p-value lower than 0.1. Ribosomal genes were filtered out and the resulting gene lists used for subsequent OAR tests.

#### OAR score associated gene expression

To identify genes associated with OAR scores we used a likelihood-ratio test implemented in GlmGamPoi^16^. Briefly, we retrieved raw count matrices from naive and activated human plasmacytoid dendritic cells (GSE157305), developing murine embryonic pancreatic epithelial cells (GSE132188), and vegetative plus starved *Dictyostelium discoideum* cells (GSE164010), then we used ‘calculateSumFactors’ from scran^37^ to estimate size factors for normalization. Finally a design matrix was built based on either the OAR score of each cell in the dataset alone, or the natural splines calculated on the OAR scores using the ‘ns’ function in splines^29,38^. Expression matrices, size factors and model matrices were then used in ‘glm_gp’ and ‘test_de’ from GlmGamPoi v. 1.16.0 to identify genes significantly associated with OAR scores compared to a reduced design where all cells had a coefficient of 1. Given the number of cells tested, test_de produces overly optimistic p-values of the predicted fits, consequently, we set the adjusted p-value threshold based on the dimensions of the data, like so,

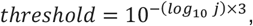

where *j* is the number of cells in the dataset. OAR associated genes are those with an adjusted p-value equal to or lower than this calculated threshold.

#### Pathway enrichment and scoring

OAR associated genes were analysed with String.DB^39^ to identify enriched pathways within Gene ontology Biological Processes and kyoto encyclopedia of genes and genomes (KEGG) pathways. Gene identifiers associated with enriched pathways were retrieved from their corresponding databases, and used to calculate expression scores with ‘AddModuleScore_UCell’ from UCell v. 2.8.0^40^ with default parameters.

#### Feature selection with OAR associated genes

OAR associated genes were used in the Seurat v.5^30^ analysis pipeline described above. Specifically, principal components were calculated using only OAR associated genes with ‘RunPCA’, setting the ‘features’ argument to only this gene set. All subsequent steps were applied as described before, so that batch correction, visualization and clustering were obtained from the 30 principal components calculated based on OAR associated genes.

#### *Dictyostelium discoideum* maintenance and analysis

*Dictyostelium discoideum* strain Ax4 was purchased from the Dictybase stock centre. Vegetatively growing cells were axenically maintained at 21°C and 160 revolution per minute in HL5 nutrient medium (14.3g/L bacto peptone, 7.15g/L yeast extract, 18g/L maltose monohydrate, 0.641g/LNa_2_HPO_4_, 0.49g/L KH_2_PO_4_, supplemented with biotin, cyanocobalamin, folic acid, lipoic acid, riboflavin and thiamine-HCl, ampicillin and streptomycin). *Dictyostelium discoideum* were cultured as above in low fluorescence axenic LoFlo medium (ForMedium) overnight before all experiments, and then maintained in LoFlo medium in place of HL5 medium for the duration of the experiments.

For flow cytometry analysis, starvation and consequent aggregation were induced by washing *D. discoideum* four times in development buffer (5mM Na_2_HPO_4_, 5mM KH_2_PO_4_, 1mM CaCl_2_, 2mM MgCl_2_ in autoclaved H_2_O) and plating at a density of 2×10^6^ cells/mL in 1mL development buffer on tissue culture-treated plates, without shaking. Vegetatively growing cells were plated in LoFlo medium at the same density. 30 min before the end of the experimental condition, cells were collected, disaggregated and stained in PBS for 30 min.

This was done to diminish any residual autofluorescence in LoFlo medium, to stop any binding of peptone or yeast extract proteins binding to cell stains, and to ensure that differences in flow cytometric results were not due to staining in different base media of different fluorescence. Cells were collected using the BD FACSymphony™ A5 Cell Analyzer (BD Biosciences), with the software FACSDiva (BD Biosciences). An example gating strategy for identifying live, single vegetative or starved *D. discoideum* cells is shown in **Extended Data Fig. 8a**. Analysis was performed using FlowJo software (TreeStar). Dyes used were MitoTracker Red and Live/Dead Aqua (all from ThermoFisher Scientific), or JC-1 (Thermo scientific).

For microscopy analysis, 2×10^6^ cells per condition/replicate were plated in 1mL development buffer supplemented with 5μg/mL JC-1 in a 12-well tissue culture-treated plate, and allowed to adhere for 1h, ensuring consistent adherence between samples. Images were taken at 10 minute intervals with a Nikon A1+ Confocal Microscope (Nikon) using a 10X objective. Volume reconstructions were made by collecting images at 20 focal planes spaced by 2.8μm gaps. Acquired images were denoised using Nikon’s proprietary software. Image analysis was carried out using Fiji^41^.

Statistical analysis was performed using Prism 10 software (GraphPad). Comparisons for two groups were calculated using unpaired two-tailed Student’s t-tests, with a Sidak correction for multiple testing. Exact p-values are indicated in the figures.

### Computational benchmarking

We quantified the time each step of the OAR test algorithm takes to generate results, using 3 publicly available datasets employed throughout the study (pDCs - GSE157305; murine embryonic pancreatic epithelial cells - GSE132188; *D. discoideum* - GSE164010), subsampling 20 times across several orders of magnitudes of cells (**Extended Data Fig. 9**). The base R function ‘system.time’ was used to monitor wall time. We found that the calculation of Hamming distances and OAR scores scaled linearly with cell number (**Extended Data Fig. 9a**,**c**), given a set number of threads specified for Hamming distance calculation using FastHamming v. 1.2^42^. MDP assignment remained relatively stable (**Extended Data Fig. 9b**) across all cell numbers examined.

## Code availability

All code to generate figures and reproduce the analysis can be found at https://github.com/Sanin-Lab/OAR-manuscript. The accompanying R package with instructions for installation and exploration of other datasets can be retrieved from https://sanin-lab.github.io/OARscRNA.

## Data availability

All data used in this study is publicly available. Accession numbers for specific datasets are listed alongside a description of the data in the supplemental information Table 1.

