## Extended Data Figures for "Missing data in single-cell transcriptomes reveals transcriptional shifts"

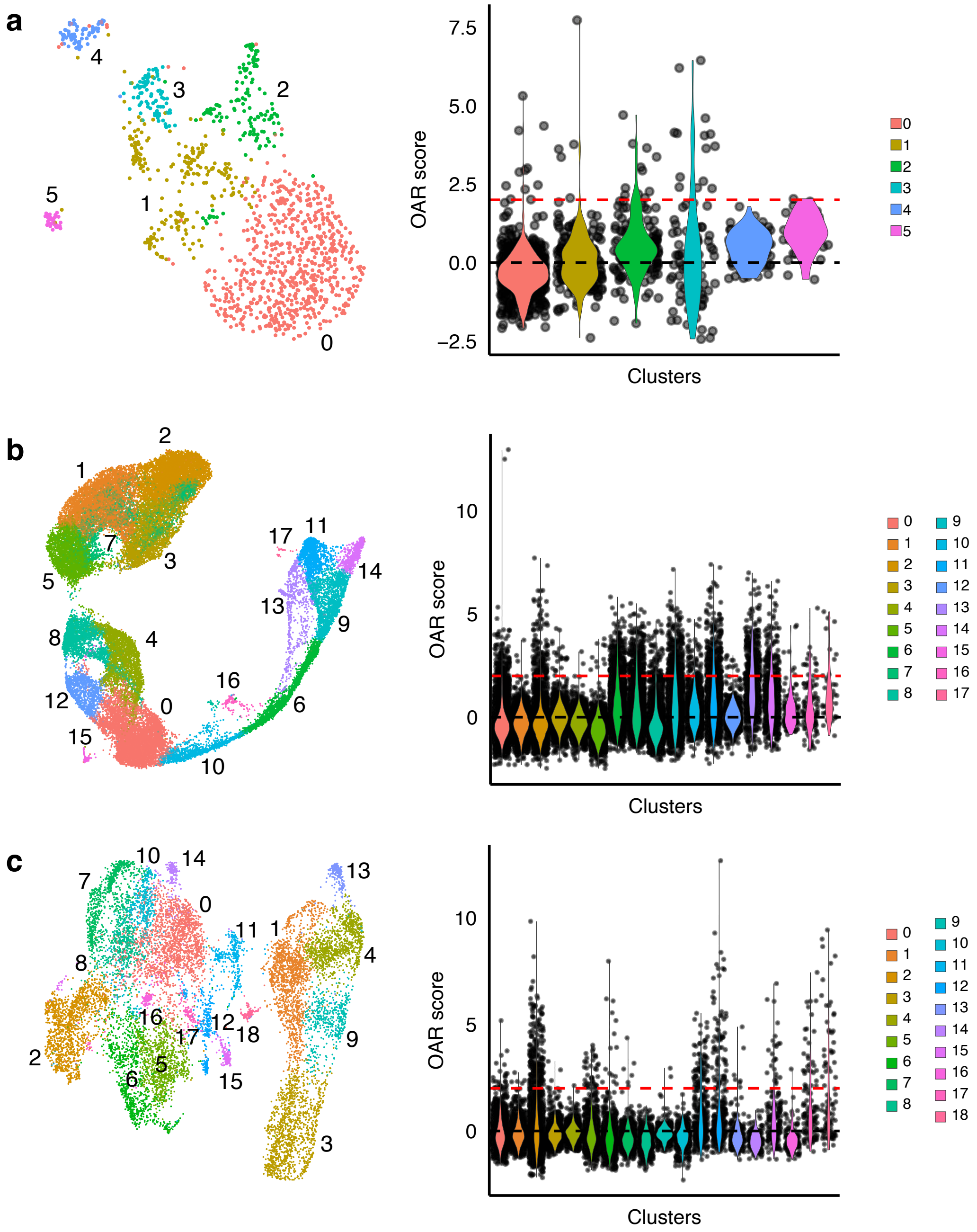
a
4
3
2
5
1
0
7.5
5.0
2.5
0.0
−2.5
OAR score
Clusters
0
1
2
3
4
5
b
2
1
7
3
5
17
11
14
13
9
8
4
16
6
12
0
15
10
10
5
0
OAR score
Clusters
c
10
14
7
0
13
11
1
4
8
16
12
18
17
9
2
6
5
15
3
10
5
0
OAR score
Clusters

***Extended data figure 1. OAR scores are unevenly distributed across cell clusters.***

**a-c**, UMAP showing cluster labels (left) and OAR scores per cluster as violin plots (right) for pDCs (**a**), murine embryonic pancreatic epithelial cells (**b**) and *D. discoideum* (**c**), with cells represented by points. Red dashed line set to OAR score of 2 (i.e. 2 standard deviations greater than the mean -$\log_{10}$ adjusted Kruskal-Wallis p-value).

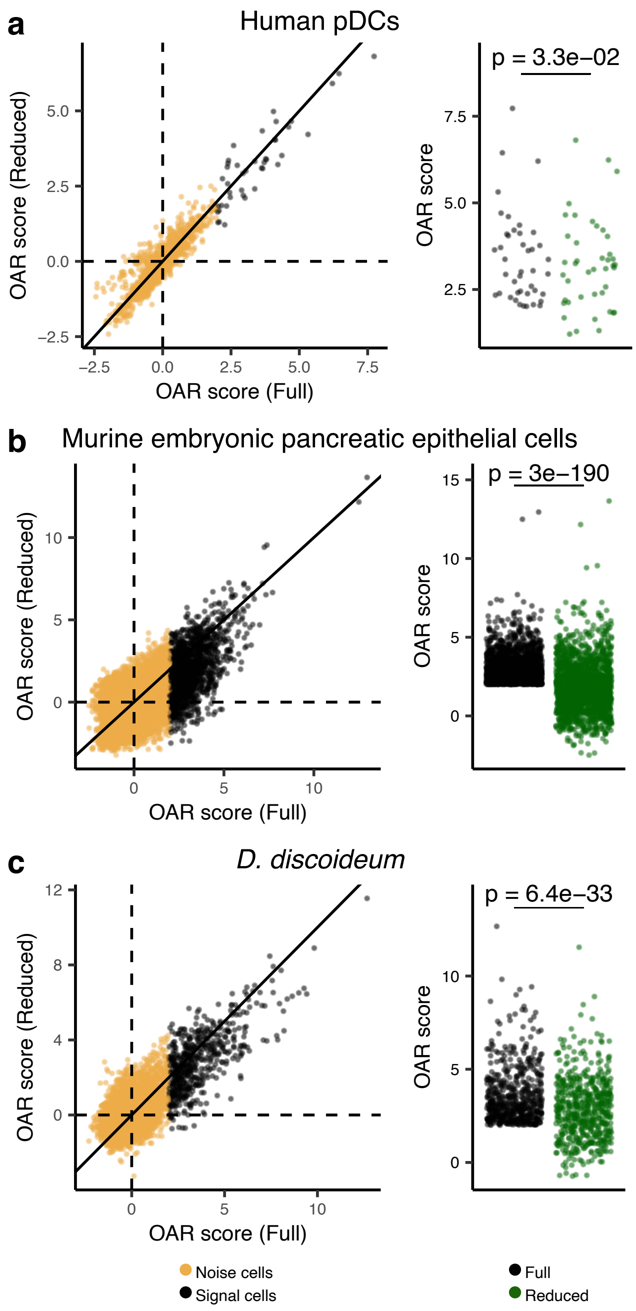
a
Human pDCs
OAR score (Reduced)
OAR score (Full)
OAR score
p = 3.3e−02
b
Murine embryonic pancreatic epithelial cells
OAR score (Reduced)
OAR score (Full)
OAR score
p = 3e−190
c
D. discoideum
OAR score (Reduced)
OAR score (Full)
OAR score
p = 6.4e−33
Noise cells
Signal cells
Full
Reduced

***Extended data figure 2. OAR scores depend on differentially expressed genes within signal cells.***

**a-c**, OAR scores were calculated for pDCs (**a**), murine embryonic pancreatic epithelial cells (**b**) and *D. discoideum* (**c**), using all genes above a set filtering threshold of 1% (full) or after removing differentially expressed genes for cells with a high OAR score (reduced). (Left) Black points represent cells with an OAR score greater than 2 in the full model (signal cells), while yellow points represent all other cells in the dataset (noise cells). Solid black line corresponding to $y = x$ is included to illustrate change in OAR scores. (Right) OAR score differences in signal cells as points in the full (black) or reduced (green) gene expression matrices are also shown. A paired Wilcoxon signed rank test was used to compare OAR scores between approaches, with p-values reported for each comparison.

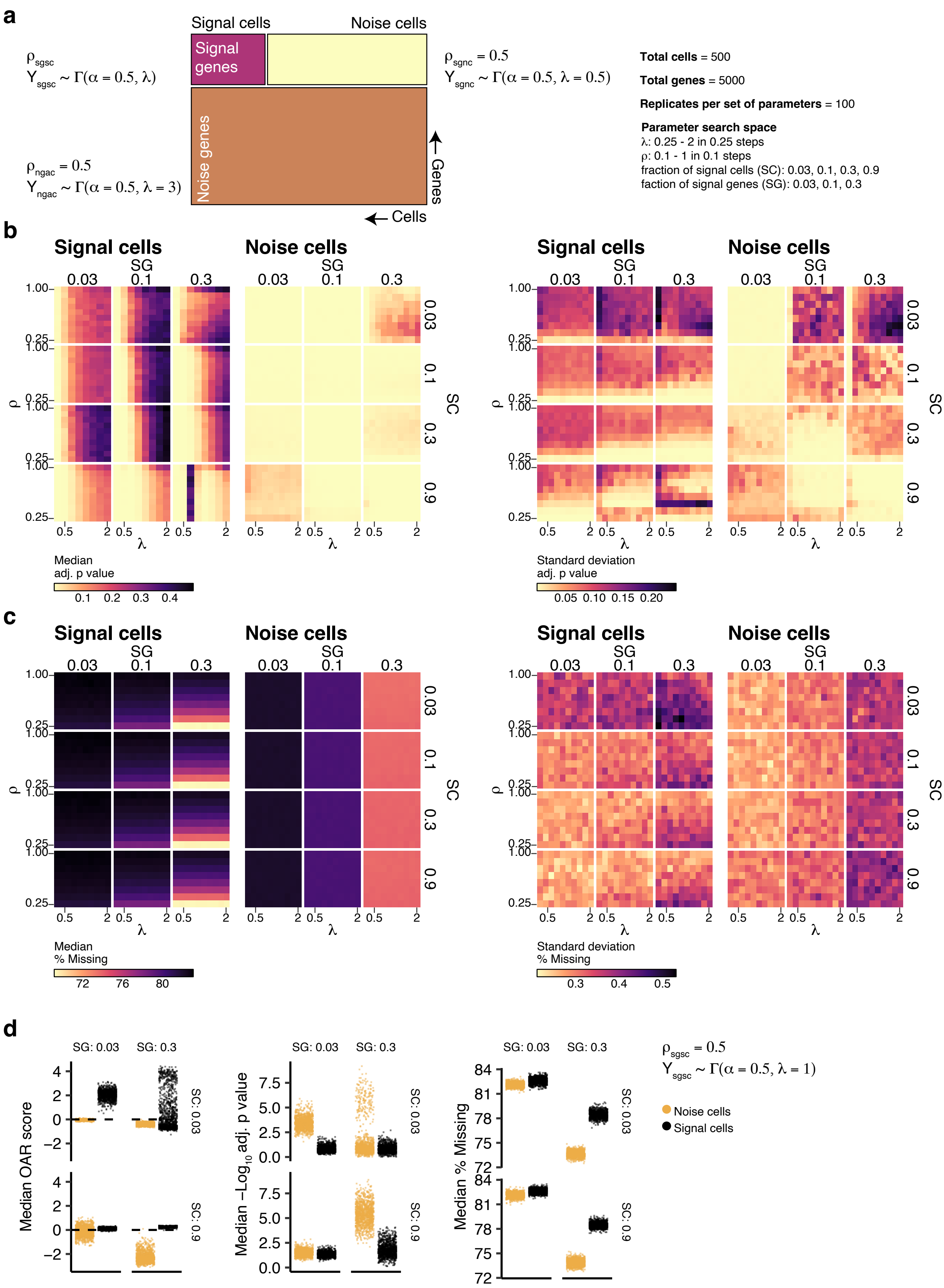

a
Signal cells
Noise cells
Signal genes
Noise genes
Genes
Cells
ρsgsc
Ysgsc ~ Γ(α = 0.5, λ)
ρsgnc = 0.5
Ysgnc ~ Γ(α = 0.5, λ = 0.5)
ρngac = 0.5
Yngac ~ Γ(α = 0.5, λ = 3)
Total cells = 500
Total genes = 5000
Replicates per set of parameters = 100
Parameter search space
λ: 0.25 - 2 in 0.25 steps
ρ: 0.1 - 1 in 0.1 steps
fraction of signal cells (SC): 0.03, 0.1, 0.3, 0.9
faction of signal genes (SG): 0.03, 0.1, 0.3
b
Signal cells
Noise cells
SG
SC
Median adj. p value
Standard deviation adj. p value
c
Median % Missing
Standard deviation % Missing
d
SG: 0.03
SG: 0.3
SC: 0.03
SC: 0.9
Median OAR score
Median −Log10 adj. p value
Median % Missing
ρsgsc = 0.5
Ysgsc ~ Γ(α = 0.5, λ = 1)
Noise cells
Signal cells

***Extended data figure 3. Simulation strategy and parameter adjustment.***

**a**, Schematic overview of simulation approach. Gene expression matrices (cells = 500, genes = 5000) were simulated using Gamma-Poisson distributions as three separate components each: i) signal cells (i.e. those with an OAR score > 2) expressing signal genes (i.e. genes responsible for signal); ii) noise cells (i.e. those with an OAR score < 2) expressing signal genes; iii) all cells expressing noise genes. Rate ($\lambda$), shape ($\alpha$) and correlation ($\rho$) for each component, or the range from which they were drawn is specified. A range of proportions of signal cells to total cells (SC) and signal genes to total genes (SG) were also examined. Simulations were repeated for each combination of parameters 100 times. **b,c**, Heatmaps show median (left) and standard deviation (right) across 100 simulations for the median adjusted p-value (**b**), and median proportion of genes with 0 counts (% Missing - **c**) calculated within each simulation for signal and noise cells, for each combination of parameters. **d**, Individual simulation results, across 1000 replications for indicated parameter combinations are also shown. Points represent individual replicates.

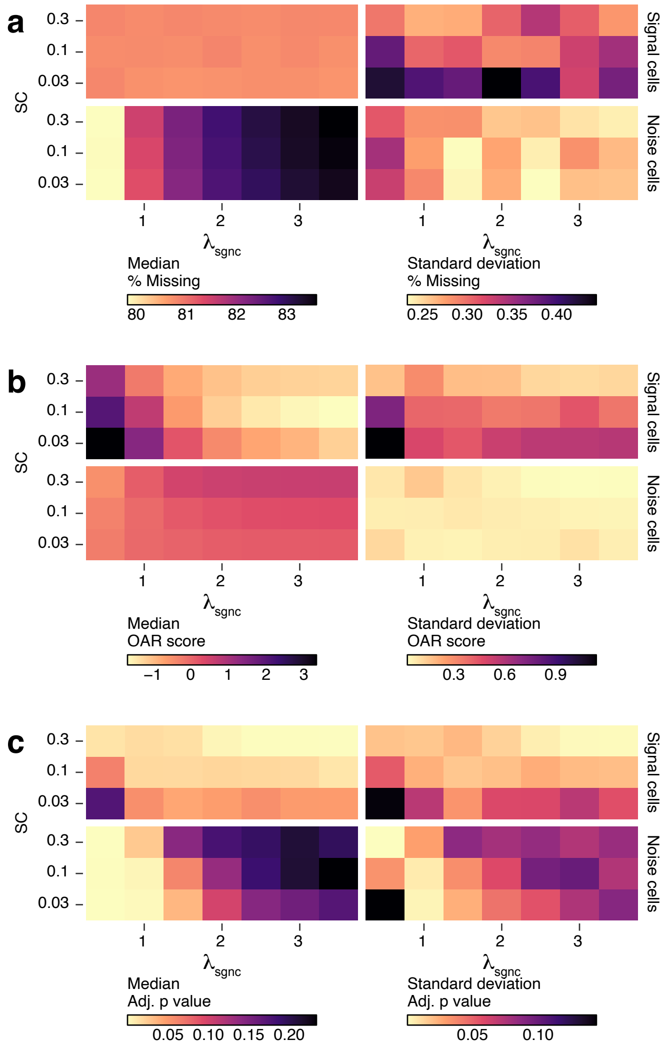

a
b
c
SC
0.3
0.1
0.03
Signal cells
Noise cells
1
2
3
$\lambda_{sgnc}$
Median
% Missing
80 81 82 83
Standard deviation
% Missing
0.25 0.30 0.35 0.40
Median
OAR score
−1 0 1 2 3
Standard deviation
OAR score
0.3 0.6 0.9
Median
Adj. p value
0.05 0.10 0.15 0.20
Standard deviation
Adj. p value
0.05 0.10

***Extended data figure 4. Simulating high sparsity in noise cells.***

**a-c**, Gene expression matrices (cells = 500, genes = 5000) were simulated using Gamma-Poisson distributions as in Extended Data Fig. 3, with $\lambda_{sgsc} = 0.75$, $\alpha_{sgsc} = 0.5$ and $\rho_{sgsc} = 0.1$, while the rate for signal genes in noise cells ($\lambda_{sgnc}$) was set as indicated. All other parameters were set as described before. Heatmaps show median (left) and standard deviation (right) across 100 simulations for the median proportion of genes with 0 counts (% Missing - **a**), the median OAR score (**b**), and the median adjusted p-value (**c**), calculated within each simulation for signal and noise cells, for each combination of parameters.

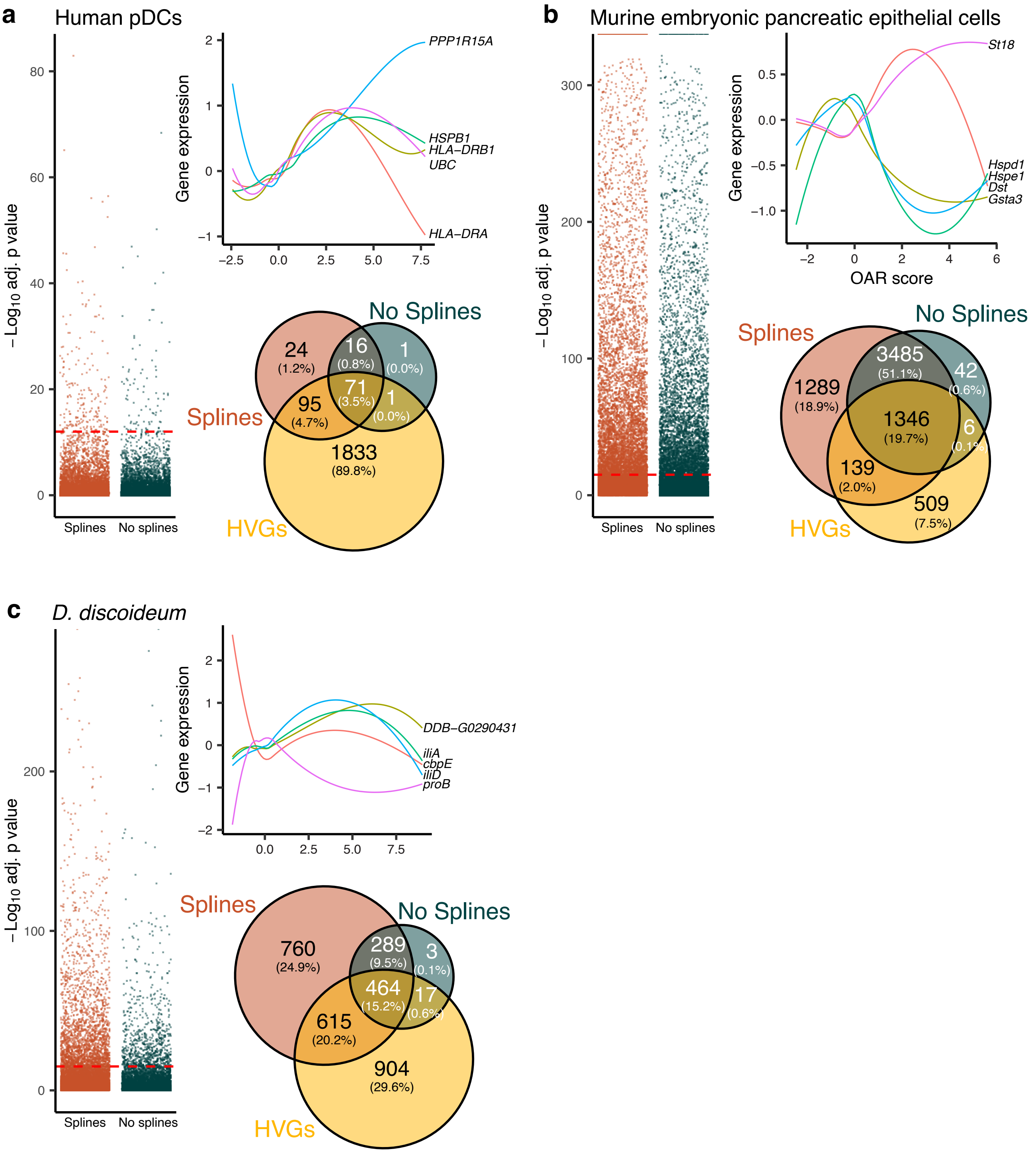
a
Human pDCs
−Log10 adj. p value
Splines
No splines
Gene expression
PPP1R15A
HSPB1
HLA−DRB1
UBC
HLA−DRA
Splines
No Splines
HVGs
24 (1.2%)
16 (0.8%)
1 (0.0%)
71 (3.5%)
95 (4.7%)
1 (0.0%)
1833 (89.8%)
b
Murine embryonic pancreatic epithelial cells
−Log10 adj. p value
Splines
No splines
Gene expression
OAR score
St18
Hspd1
Hspe1
Dst
Gsta3
Splines
No Splines
HVGs
1289 (18.9%)
3485 (51.1%)
42 (0.6%)
1346 (19.7%)
6 (0.1%)
139 (2.0%)
509 (7.5%)
c
D. discoideum
−Log10 adj. p value
Splines
No splines
Gene expression
DDB−G0290431
iliA
cbpE
iliD
proB
Splines
No Splines
HVGs
760 (24.9%)
289 (9.5%)
3 (0.1%)
464 (15.2%)
17 (0.6%)
615 (20.2%)
904 (29.6%)

***Extended data figure 5. Identifying OAR associated genes using likelihood ratio test with splines.***

**a-c**, A likelihood ratio test was used to identify genes significantly associated with OAR scores across all cells in indicated datasets. Model matrices using either OAR scores or OAR score-calculated natural splines to capture non-linear relationships were employed, testing against a reduced model (~1). Adjusted p-values for the fit of each morel across all genes are shown (left), with 5 most significant results using natural splines presented (top right) alongside the overlap between both methods and highly variable features (HVGs) obtained using FindVariableFeatures (bottom right). Red dashed lines indicate the adjusted p-value threshold used in the analysis to consider a fit to be significant.

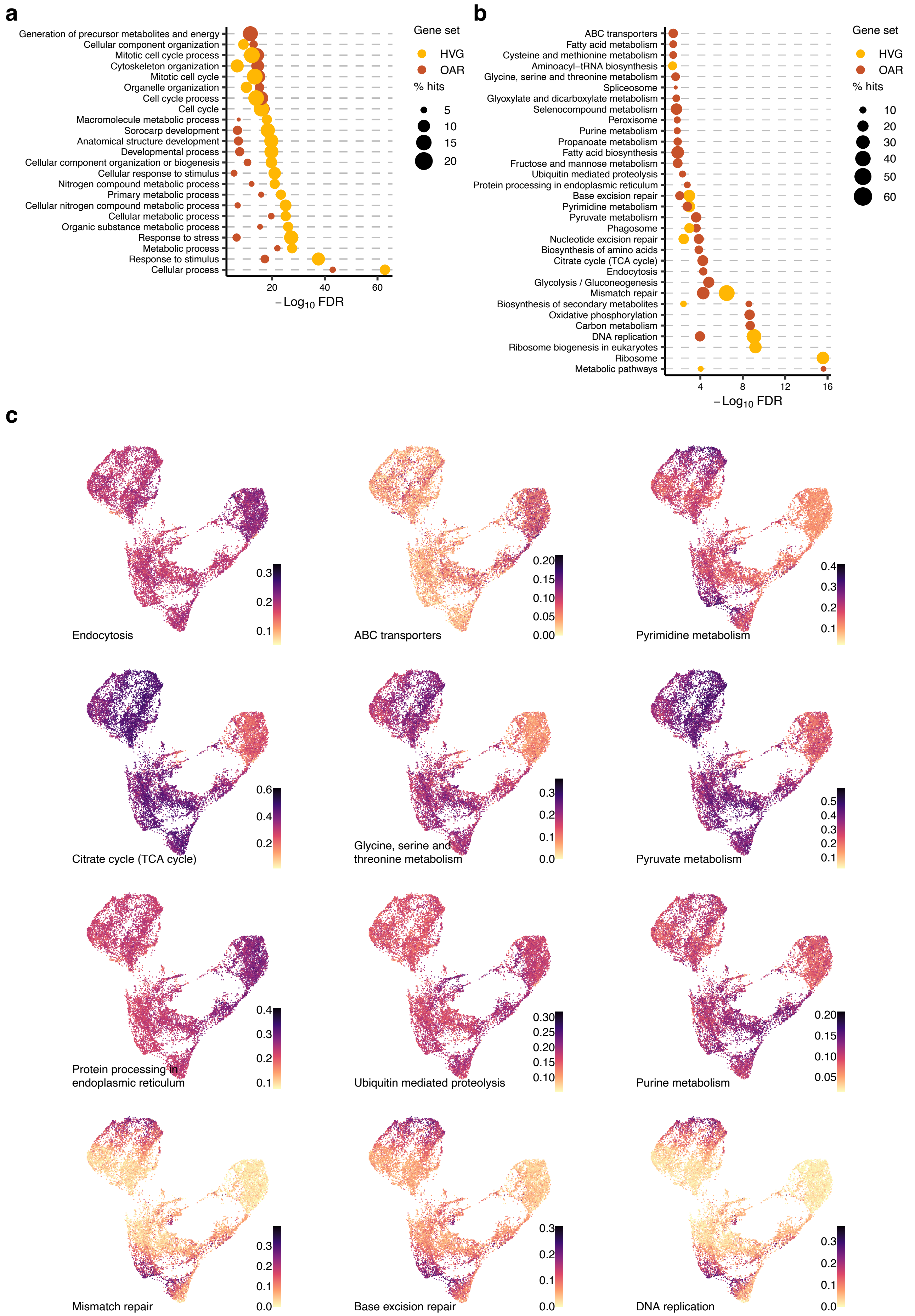
a
Generation of precursor metabolites and energy
Cellular component organization
Mitotic cell cycle process
Cytoskeleton organization
Mitotic cell cycle
Organelle organization
Cell cycle process
Cell cycle
Macromolecule metabolic process
Sorocarp development
Anatomical structure development
Developmental process
Cellular component organization or biogenesis
Cellular response to stimulus
Nitrogen compound metabolic process
Primary metabolic process
Cellular nitrogen compound metabolic process
Cellular metabolic process
Organic substance metabolic process
Response to stress
Metabolic process
Response to stimulus
Cellular process
20
40
60
$-\text{Log}_{10}$ FDR
Gene set
HVG
OAR
% hits
5
10
15
20
b
ABC transporters
Fatty acid metabolism
Cysteine and methionine metabolism
Aminoacyl−tRNA biosynthesis
Glycine, serine and threonine metabolism
Spliceosome
Glyoxylate and dicarboxylate metabolism
Selenocompound metabolism
Peroxisome
Purine metabolism
Propanoate metabolism
Fatty acid biosynthesis
Fructose and mannose metabolism
Ubiquitin mediated proteolysis
Protein processing in endoplasmic reticulum
Base excision repair
Pyrimidine metabolism
Pyruvate metabolism
Phagosome
Nucleotide excision repair
Biosynthesis of amino acids
Citrate cycle (TCA cycle)
Endocytosis
Glycolysis / Gluconeogenesis
Mismatch repair
Biosynthesis of secondary metabolites
Oxidative phosphorylation
Carbon metabolism
DNA replication
Ribosome biogenesis in eukaryotes
Ribosome
Metabolic pathways
4
8
12
16
$-\text{Log}_{10}$ FDR
Gene set
HVG
OAR
% hits
10
20
30
40
50
60
c
Endocytosis
ABC transporters
Pyrimidine metabolism
Citrate cycle (TCA cycle)
Glycine, serine and threonine metabolism
Pyruvate metabolism
Protein processing in endoplasmic reticulum
Ubiquitin mediated proteolysis
Purine metabolism
Mismatch repair
Base excision repair
DNA replication

***Extended data figure 6. Pathway enrichment analysis and scoring in D. discoideum.***

**a,b**, Pathway enrichment analysis results for Highly Variable Genes (HVG - yellow) and OAR associated genes (brown) against Gene ontology Biological Processes (**a**) or KEGG pathways (**b**). **c**, UMAPs with cells colored by indicated enriched pathway scores.

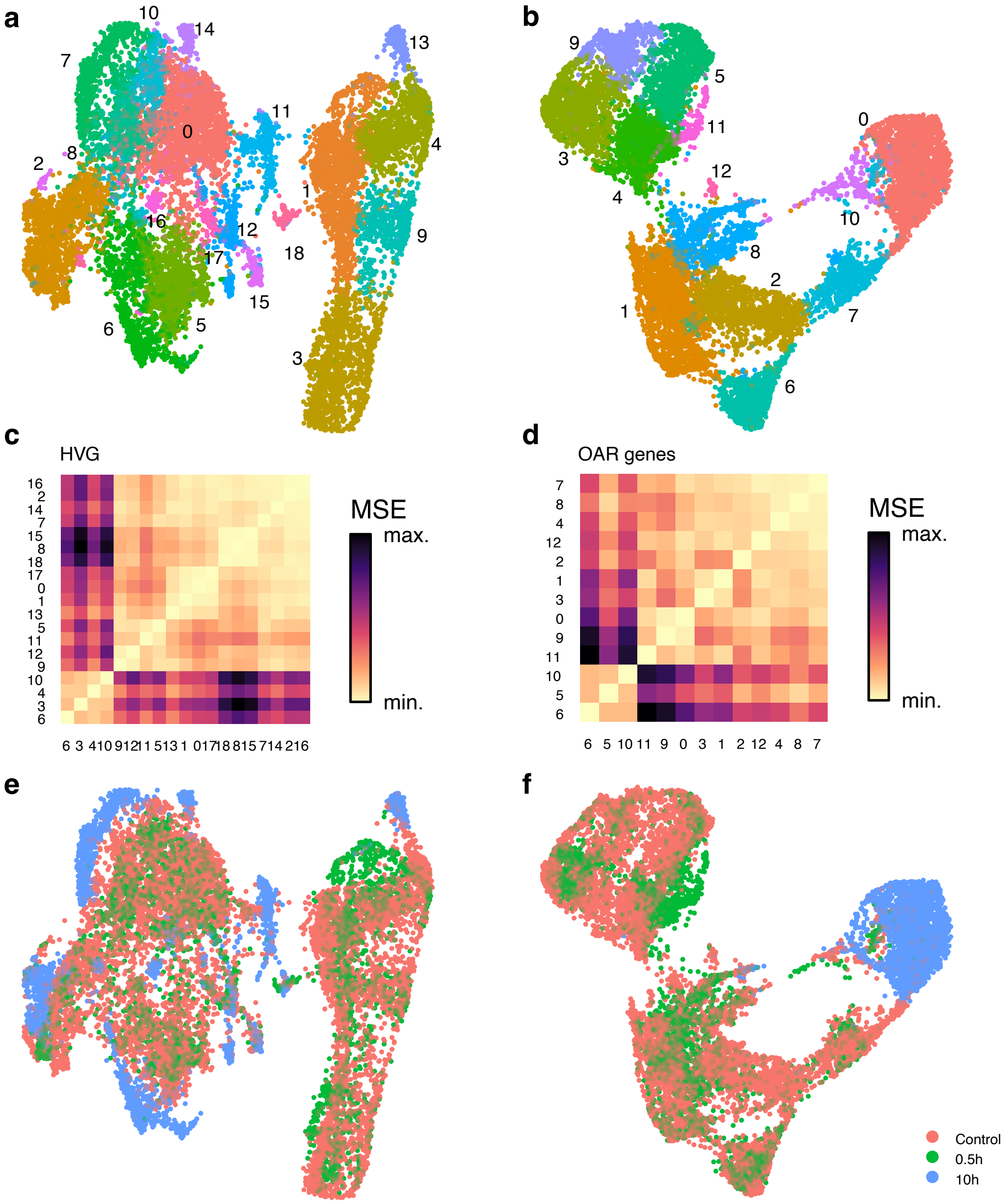
a
b
c
HVG
MSE
max.
min.
d
OAR genes
MSE
max.
min.
e
f
Control
0.5h
10h

***Extended data figure 7. Dimensionality reduction and clustering using OAR associated genes.***

**a-f**, *D. discoideum* UMAPs with labelled clusters (**a,b**), mean standard errors between clusters (**c,d**) and UMAPs labelled according to experimental groups (**e,f**) are shown. Clustering and dimensionality reduction were calculated based on highly variable genes (**a**, **c**, **e**) or OAR associated genes (**b**, **d**, **f**).

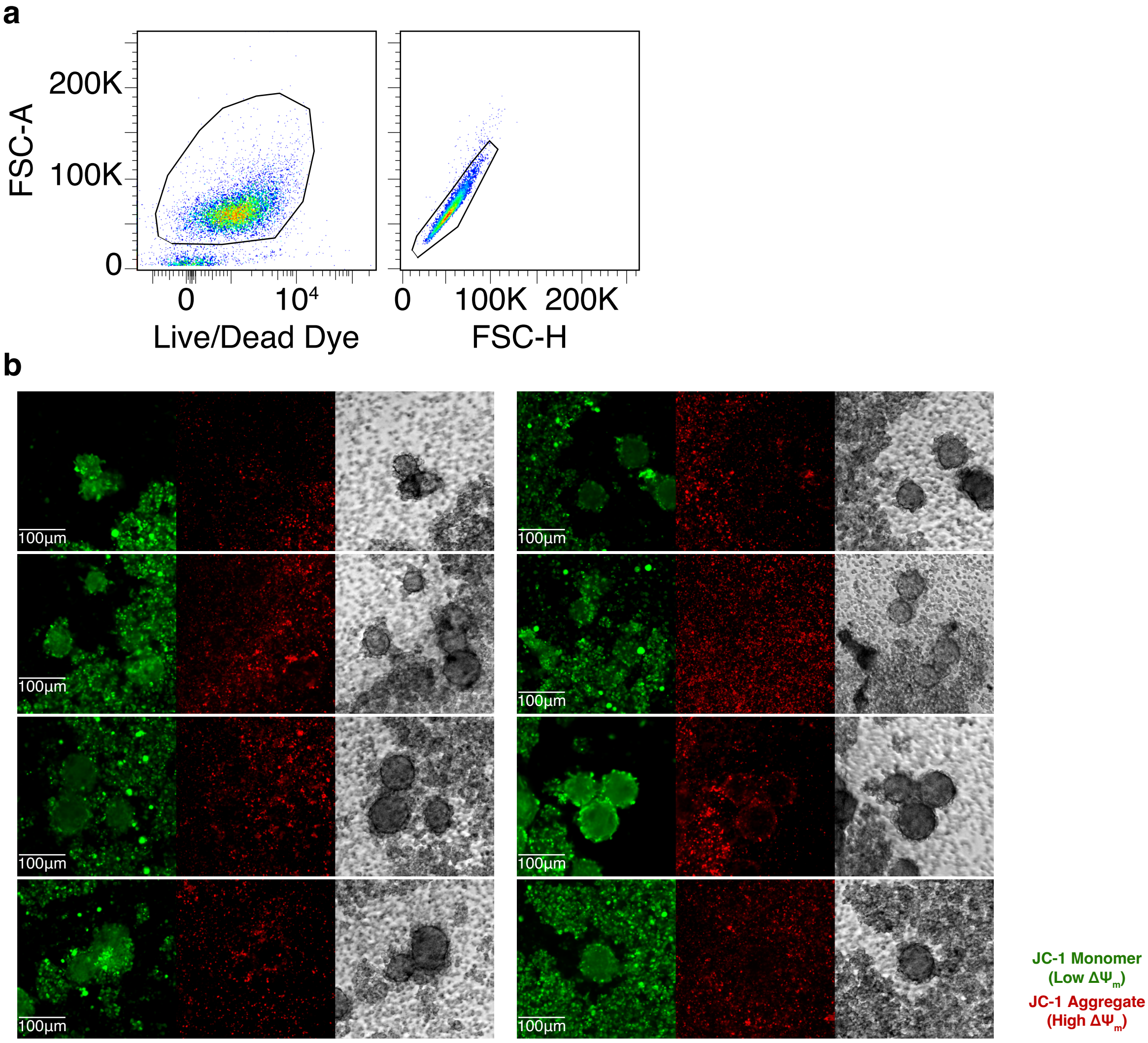
a
FSC-A
200K
100K
0
0
10⁴
Live/Dead Dye
0
100K
200K
FSC-H
b
100µm
JC-1 Monomer
(Low ΔΨm)
JC-1 Aggregate
(High ΔΨm)

***Extended data figure 8. D. discoideum cells decrease mitochondrial activity as they form multicellular aggregates.***

**a**, Gating strategy for *D. discoideum* cells. **b**, *D. discoideum* cells were starved for 24h then examined for mitochondrial activity with mitochondrial membrane potential ($\Delta\Psi_m$) sensing probe JC-1 via microscopy. Cells forming multicellular aggregates had consistently depolarized mitochondria, evident as JC-1 monomers (510nm excitation / 527nm emission), whereas cells in the periphery displayed high $\Delta\Psi_m$, evident as JC-1 aggregates (585nm excitation / 590nm emission). Representative images from 2 independent experiments are shown.

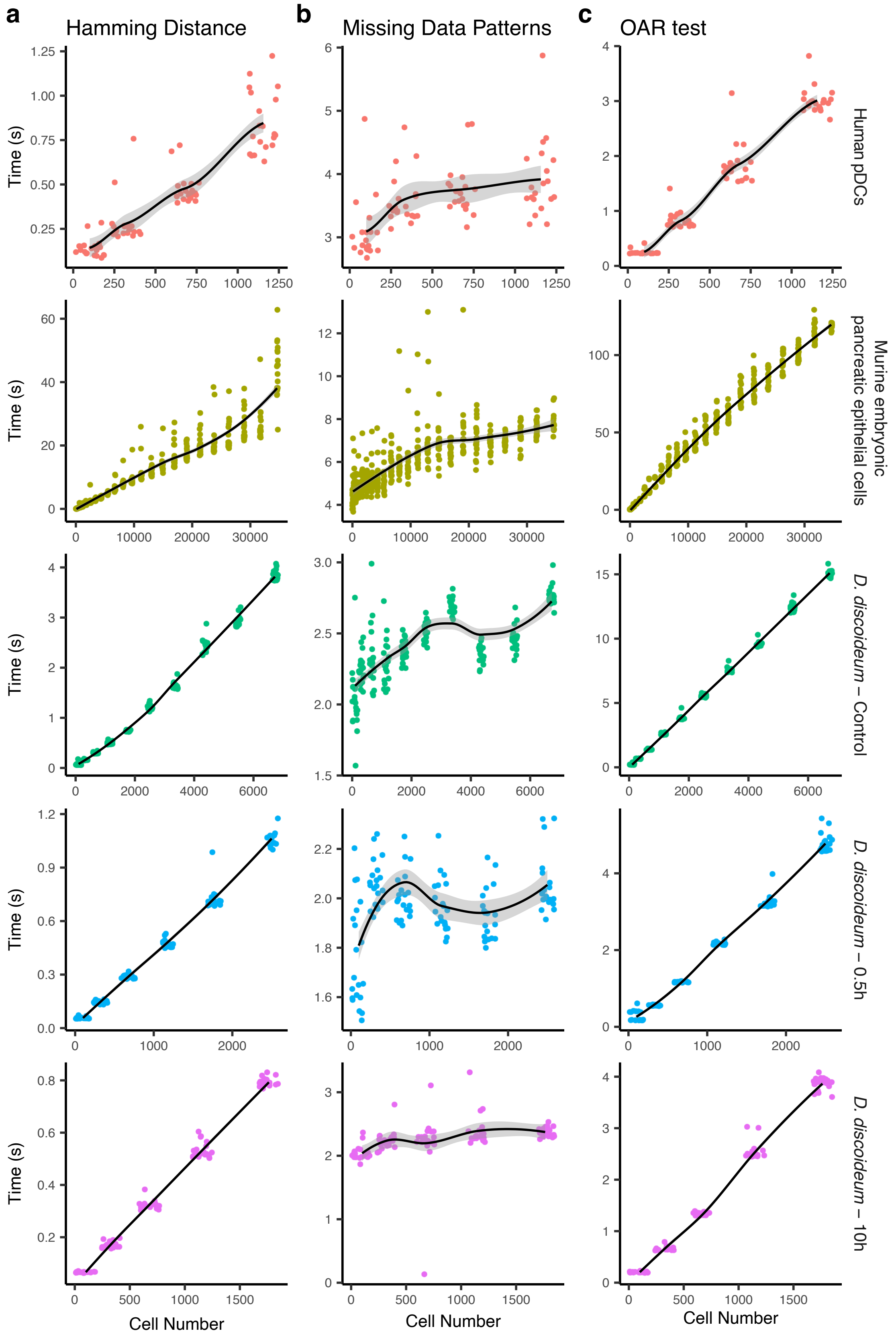
a
Hamming Distance
b
Missing Data Patterns
c
OAR test
Human pDCs
Murine embryonic pancreatic epithelial cells
D. discoideum – Control
D. discoideum – 0.5h
D. discoideum – 10h
Time (s)
Cell Number

***Extended data figure 9. Benchmark functions in OAR score calculation.***

**a-c**, Indicated datasets were subsampled iteratively (n = 20) to obtain increasing number of cells spanning the range of each dataset, then the time to complete three main steps in the OAR test algorithm, bin gene expression and calculate hamming distances (**a**), identify missing data patterns (**b**), and run Kruskal-Wallis test for each cell (**c**), were measured for each iteration independently. Points represent independent sampling iterations for indicated cell numbers. Lines represent Loess regressions with shaded areas showing standard errors.
