## Supplemental Information for "Missing data in single-cell transcriptomes reveals transcriptional shifts"

### Title

**Supplementary Table S1 - Dataset description and source**

A brief description of single cell RNA-seq datasets used in the study alongside accession codes.

| Accession | Source | Cell type | Conditions | Technology | Ref. |
|---|---|---|---|---|---|
| GSE157305 | *Homo sapiens* Blood | Plamacytoid Dendritic cells | Control or ODN activated | 10X Chromium controller | 1 |
| GSE164010 | *Dictyostelium discoideum* | - | Vegetative or Starved | 10X Chromium controller | 2 |
| GSE132188 | *Mus musculus* Pancreas | Epithelial cells | Embryonic development | 10X Chromium controller | 3 |
| GSE176078 | *Homo sapiens* Breast cancer | All cells in tissue | Breast cancer biopsies | 10X Chromium controller | 4 |
| GSE84133 | *Homo sapiens* Pancreas | All cells in tissue | Pancreas biopsies | inDrop | 5 |
| GSE85241 | *Homo sapiens* Pancreas | All cells in tissue | Pancreas biopsies | CEL-seq2 | 6 |
| E-MTAB-5061 | *Homo sapiens* Pancreas | All cells in tissue | Pancreas biopsies | Smart-Seq2 | 7 |
| GSE141115 | *Mus musculus* Kidney | All cells in tissue | Dissociation techniques | 10X Chromium controller | 8 |
| GSE171328 | *Mus musculus* Popliteal adipose | Stromal vascular fraction | Naïve control or *L. monocytogenes* infection | 10X Chromium controller | 9 |
| GSE157313 | *Mus musculus* Mesenteric adipose | Stromal vascular fraction | Naïve control or *H. polygyrus* infection | 10X Chromium controller | 10 |
| GSE171330 | *Mus musculus* Lamina propria | CD45+ cells | Control (chow) diet or High fat diet | 10X Chromium controller | 11 |
| GSE144707 | *Mus musculus* Sciatic nerve | Macrophages | Naïve control or Nerve crush | 10X Chromium controller | 12 |
| GSE152674 | *Mus musculus* Breast tumor | Myeloid cells | Spontaneous tumor in mice with WT or *Dab2* deficient macrophages | 10X Chromium controller | 13 |
| GSE123587 | *Mus musculus* Atherosclerotic plaque | Macrophages | Progressing or Regressing lesion | 10X Chromium controller | 14 |
| GSE146233 | *Mus musculus* Lung | CD45+ cells | Naïve control or *Cryptococcus neoformans* infection | 10X Chromium controller | 15 |
| GSE145086 | *Mus musculus* Liver | Non-parenchymal liver cells | Healthy control or Fibrotic tissue | 10X Chromium controller | 16 |
| GSE106472 | *Mus musculus* Heart | CD45+ cells | Healthy control or Infarcted tissue | inDrop | 17 |
| GSE121081 | *Mus musculus* Retina | CD45+ cells | Healthy (dark) control or Neurodegeneration (light) | 10X Chromium controller | 18 |

| | | | | | |
|---|---|---|---|---|---|
| GSE113111 | *Mus musculus*<br>Skeletal muscle | CD45+ cells | Naïve control or *Toxoplasma gondii* infection | 10X Chromium controller | [19] |
| GSE121654 | *Mus musculus*<br>Brain | Microglia | Steady state | 10X Chromium controller | [20] |
| GSE183489 | *Mus musculus*<br>Skin | Macrophages | Wounded skin | Smart-Seq2 | [21] |

### Supplementary Figures

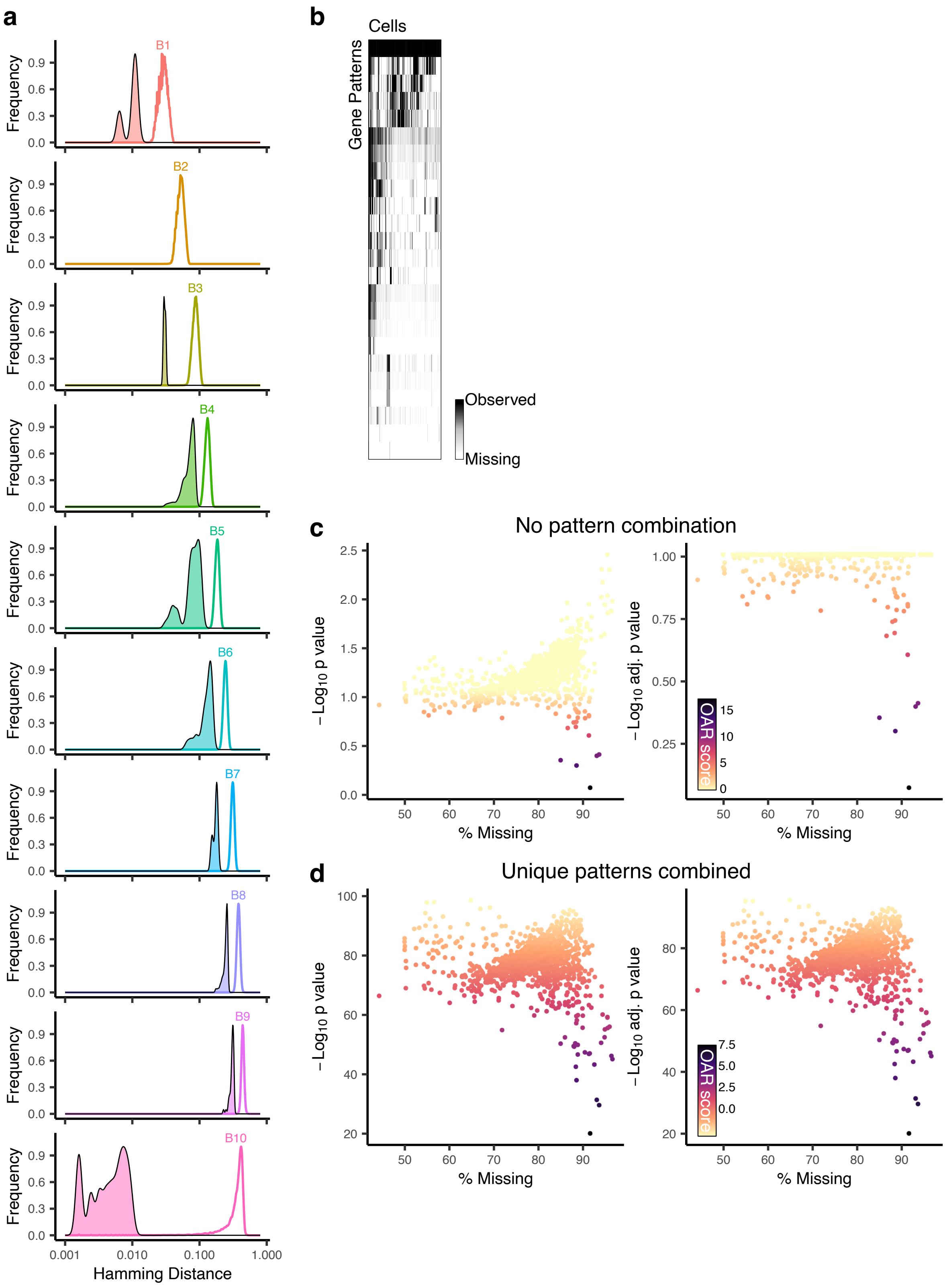
a
b
Cells
Gene Patterns
Observed
Missing
B1
B2
B3
B4
B5
B6
B7
B8
B9
B10
Frequency
Hamming Distance
0.001
0.010
0.100
1.000
c
No pattern combination
−Log10 p value
−Log10 adj. p value
% Missing
OAR score
d
Unique patterns combined
−Log10 p value
−Log10 adj. p value
% Missing
OAR score

***Supplementary Figure 1. Dynamic tolerance, pattern detection and Kruskal-Wallis results in plasmacytoid dendritic cells***

A gene expression matrix from single cell RNA-sequencing data generated from resting or activated plasmacytoid dendritic cells (pDCs) was processed and genes were separated into 10 bins according to the fraction of cells in which they were observed. Hamming distances between each pair of genes within each bin were then calculated to identify missing data patterns and contrast gene expression distribution across these patterns. **a**, Histograms show the distribution of Hamming distances within each bin for all genes (open) or those with a Hamming distance below the dynamically set threshold (closed). **b**, Identified patterns (rows) containing more than 1 gene across all cells (columns) are shown as a heatmap shaded based on the fraction of genes in the pattern observed for each cell. **c,d**, Kruskal-Wallis results before (left) or after (right) Benjamini-Hochberg correction are shown for all cells (points), shaded according to calculated OAR scores, with indicated range of values. The percentage of genes with 0 counts (% Missing) per cell is also shown. Distribution tests for all patterns, including those made out of a single gene (**c**), or where ungrouped genes are combined into a single pattern (**d**), are presented.

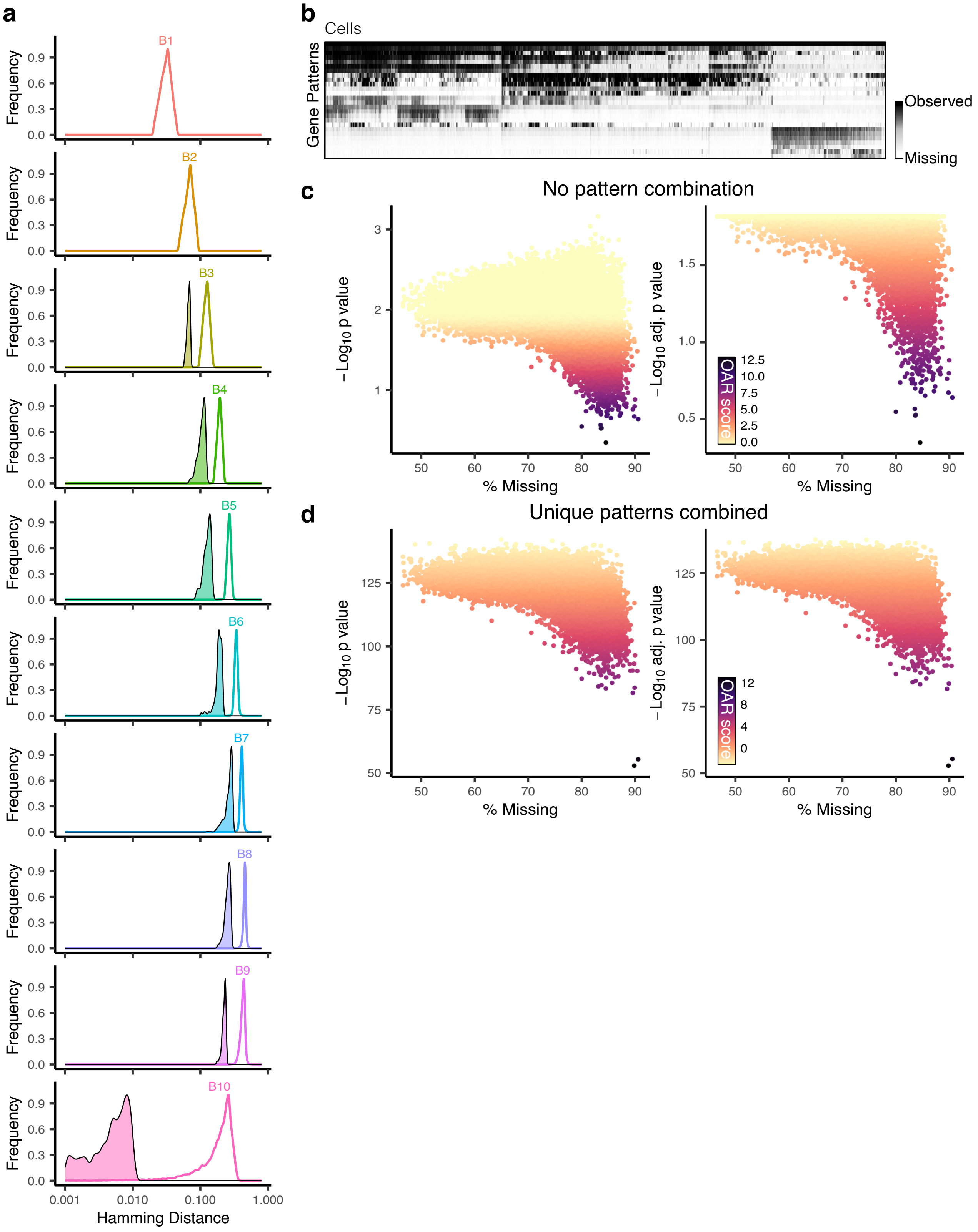
a
b
c
d
Frequency
Hamming Distance
B1
B2
B3
B4
B5
B6
B7
B8
B9
B10
0.001
0.010
0.100
1.000
Cells
Gene Patterns
Observed
Missing
No pattern combination
Unique patterns combined
−Log10 p value
−Log10 adj. p value
% Missing
OAR score

***Supplementary Figure 2. Dynamic tolerance, pattern detection and Kruskal-Wallis results in developing murine embryonic pancreatic epithelial cells.***

**a-d**, As in Supplementary Fig. 1, but applied to a gene expression matrix from single cell RNA-sequencing data generated from murine pancreatic epithelial cells obtained at different stages of embryonic development.

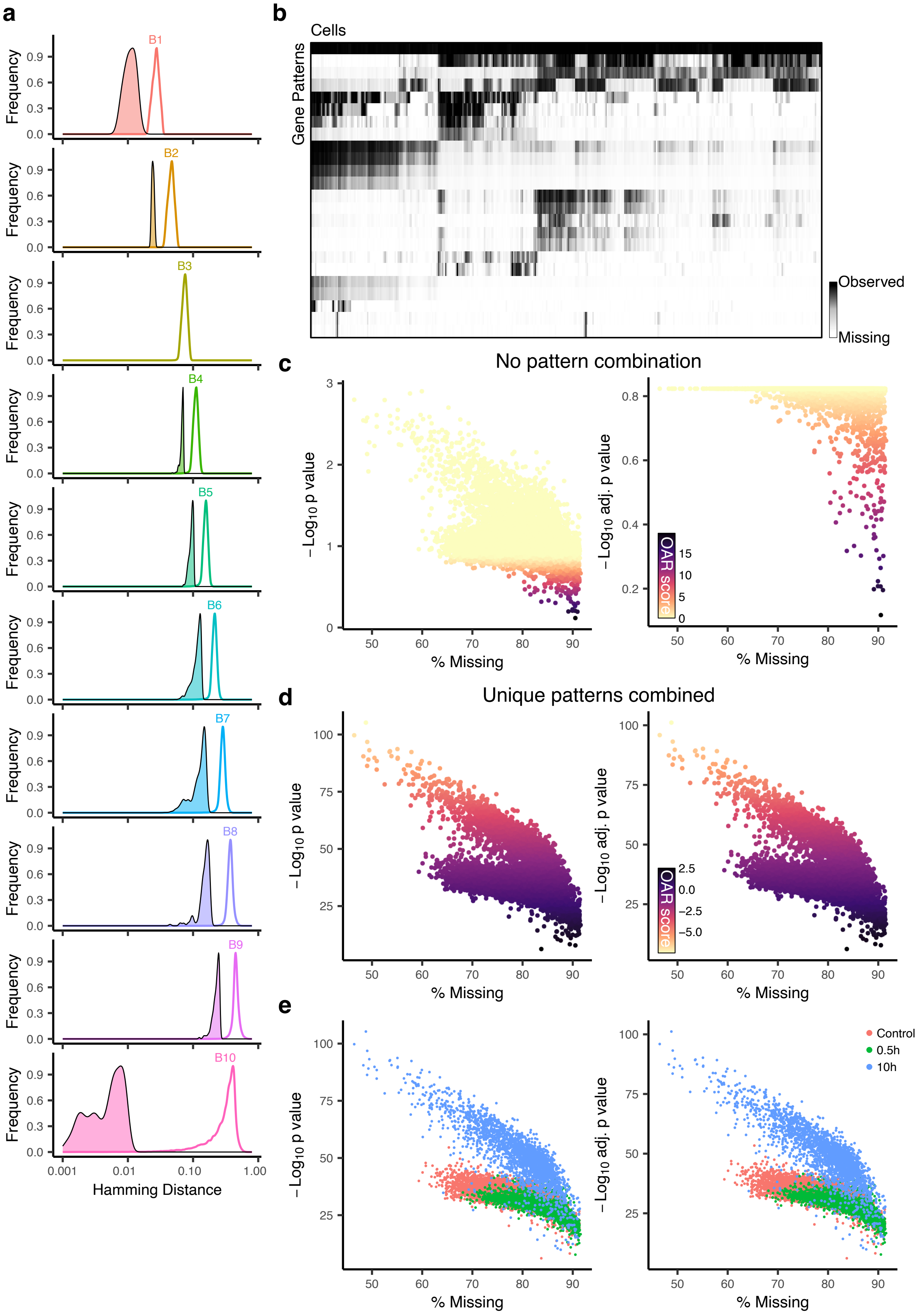

a
B1
B2
B3
B4
B5
B6
B7
B8
B9
B10
Frequency
0.0
0.3
0.6
0.9
0.001
0.01
0.10
1.00
Hamming Distance
b
Cells
Gene Patterns
Observed
Missing
c
No pattern combination
$-\mathrm{Log}_{10}$ p value
$-\mathrm{Log}_{10}$ adj. p value
% Missing
OAR score
15
10
5
0
d
Unique patterns combined
$-\mathrm{Log}_{10}$ p value
$-\mathrm{Log}_{10}$ adj. p value
% Missing
OAR score
2.5
0.0
−2.5
−5.0
e
$-\mathrm{Log}_{10}$ p value
$-\mathrm{Log}_{10}$ adj. p value
% Missing
Control
0.5h
10h

***Supplementary Figure 3. Dynamic tolerance, pattern detection and Kruskal-Wallis results in D. discoideum.***

**a-d**, As in Supplementary Fig. 1, but applied to a gene expression matrix from single cell RNA-sequencing data generated from vegetative or starved *D. discoideum* at 2 timepoints post nutrient removal. **e**, Kruskal-Wallis results before (left) or after (right) Benjamini-Hochberg correction are shown for all cells (points), shaded according to experimental groups.

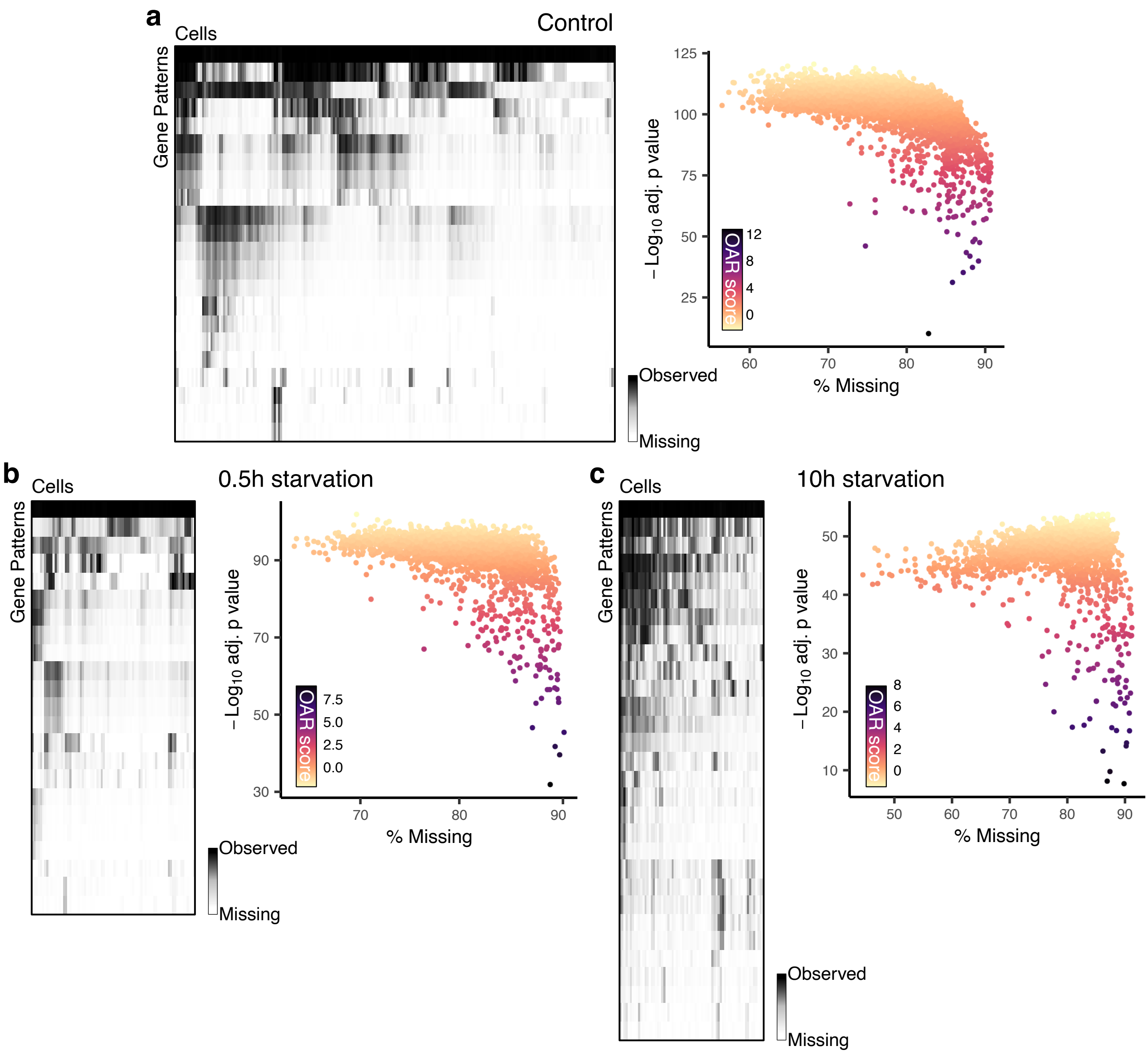

a
Cells
Control
Gene Patterns
Observed
Missing
125
100
75
50
25
−Log10 adj. p value
OAR score
12
8
4
0
60
70
80
90
% Missing
b
Cells
0.5h starvation
Gene Patterns
Observed
Missing
90
70
50
30
−Log10 adj. p value
OAR score
7.5
5.0
2.5
0.0
70
80
90
% Missing
c
Cells
10h starvation
Gene Patterns
Observed
Missing
50
40
30
20
10
−Log10 adj. p value
OAR score
8
6
4
2
0
50
60
70
80
90
% Missing

***Supplementary Figure 4. Pattern detection and Kruskal-Wallis results in D. discoideum separated by experimental group.***

**a-c**, As in Supplementary Fig. 3, but applied independently to *D. discoideum* gene expression matrices from control (**a**), 0.5h (**b**) and 10h (**c**) of starvation. Identified patterns (rows) containing more than 1 gene across all cells (columns) are shown as a heatmap shaded based on the fraction of genes in the pattern observed for each cell (left), and Kruskal-Wallis results after Benjamini-Hochberg correction are shown for all cells (points) in each experimental group (right). Points are shaded according to calculated OAR scores as indicated. The percentage of genes with 0 counts (% Missing) per cell is also shown.

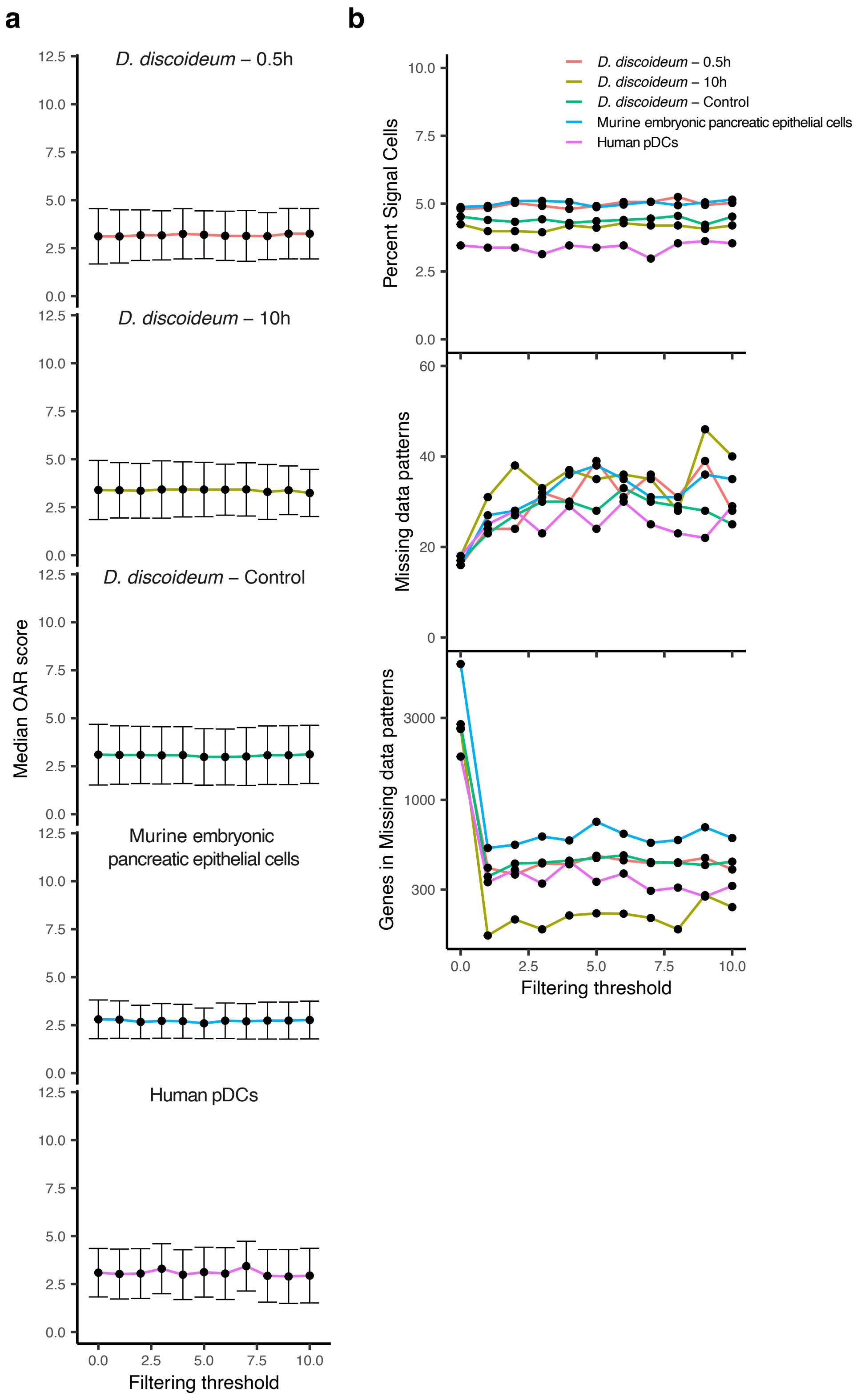
a
b
D. discoideum – 0.5h
D. discoideum – 10h
D. discoideum – Control
Murine embryonic pancreatic epithelial cells
Human pDCs
Median OAR score
Filtering threshold
Percent Signal Cells
Missing data patterns
Genes in Missing data patterns
Filtering threshold
D. discoideum – 0.5h
D. discoideum – 10h
D. discoideum – Control
Murine embryonic pancreatic epithelial cells
Human pDCs

***Supplementary Figure 5. OAR score calculations are largely independent of genes with low abundance.***

Gene expression matrices from indicated datasets were used to calculate OAR scores, and missing data patterns across filtering thresholds for gene expression based on the percentage of cells expressing a gene. Expressed genes were defined as those with a greater than 0 read count for any given cell. Signal cells were defined as those with an OAR score greater than 2 (i.e. 2 standard deviations lower than the mean-scaled $-\log_{10}$ adjusted Kruskal-Wallis p-value). **a,b**, Median OAR scores (point) and corresponding standard deviation for (**a**), and the percentage of (**b** - top), signal cells are shown across filtering thresholds. The number of detected missing data patterns and genes captured in these patterns (**b** - middle and bottom) are also shown.

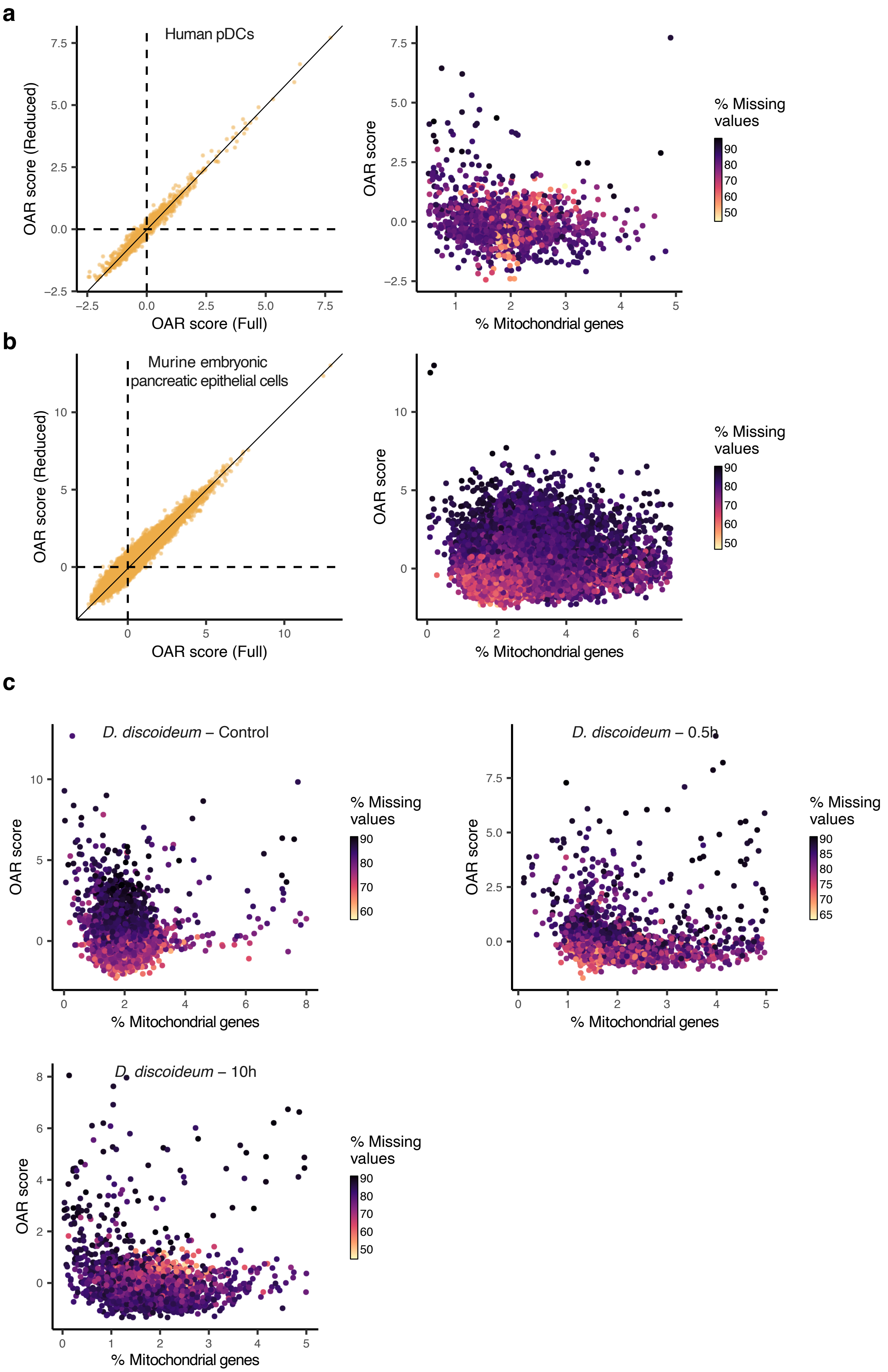
a
Human pDCs
OAR score (Reduced)
OAR score (Full)
7.5
5.0
2.5
0.0
−2.5
OAR score
% Mitochondrial genes
1
2
3
4
5
% Missing values
90
80
70
60
50
b
Murine embryonic pancreatic epithelial cells
OAR score (Reduced)
OAR score (Full)
10
5
0
OAR score
% Mitochondrial genes
2
4
6
% Missing values
90
80
70
60
50
c
D. discoideum – Control
OAR score
% Mitochondrial genes
0
2
4
6
8
10
5
% Missing values
90
80
70
60
D. discoideum – 0.5h
OAR score
% Mitochondrial genes
7.5
5.0
2.5
0.0
% Missing values
90
85
80
75
70
65
D. discoideum – 10h
OAR score
% Mitochondrial genes
8
6
4
2
0
% Missing values
90
80
70
60
50

***Supplementary Figure 6. OAR scores are independent of cell quality.***

**a,b**, OAR scores were calculated on cells based on all expressed genes above a 1% filtering threshold (full) or with blacklisted genes removed from the expression matrix prior to filtering (reduced). Blacklisted genes were identified based on SignatuR[22] and excluded from the expression matrix entirely. Results for indicated datasets are shown, with cells represented as points (left). Solid black line corresponding to $y = x$ is included to illustrate change in OAR scores. **a-c**, OAR scores and percent mitochondrial gene expression is shown for all cells (represented as points), shaded according to the percentage of missing values for indicated datasets/experimental groups.

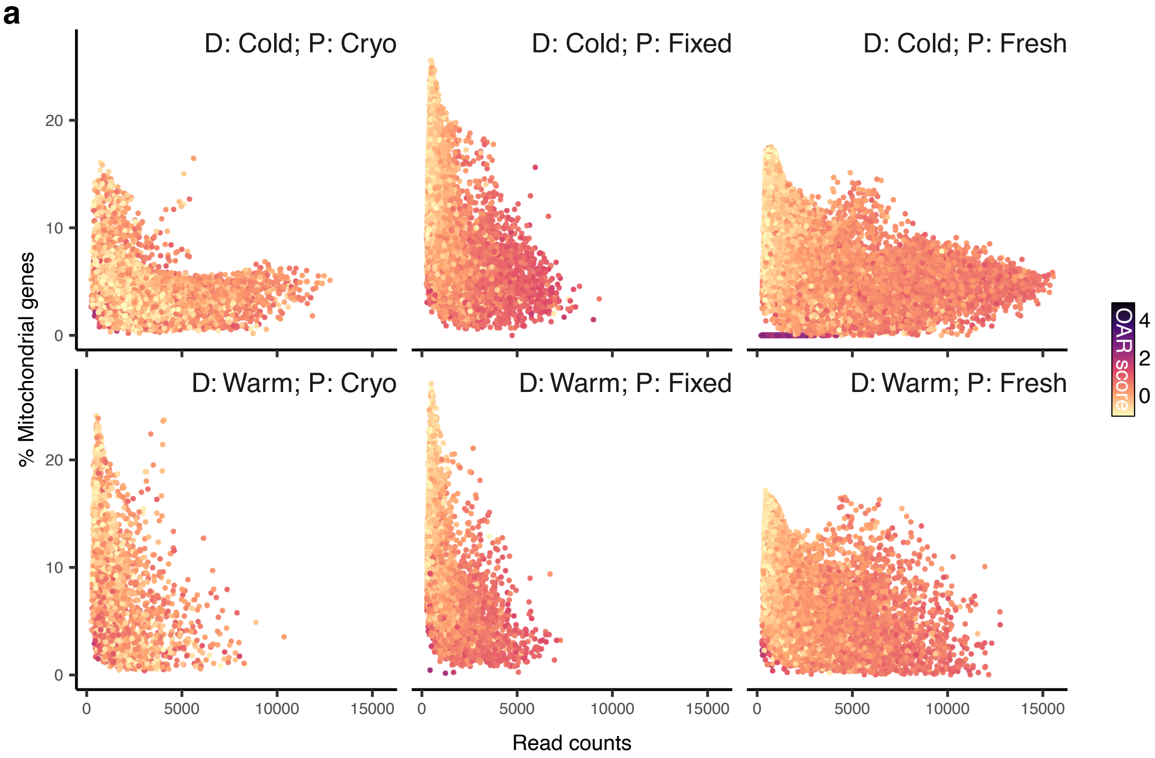
a
D: Cold; P: Cryo
D: Cold; P: Fixed
D: Cold; P: Fresh
D: Warm; P: Cryo
D: Warm; P: Fixed
D: Warm; P: Fresh
% Mitochondrial genes
Read counts
0
10
20
0
5000
10000
15000
OAR score
4
2
0

***Supplementary Figure 7. OAR scores are independent of cell preparation methods.*** OAR scores were calculated on kidney cells dissociated with 6 different tissue preparations previously shown to affect cell quality and gene expression[8]. The relationship between percent mitochondrial genes expression, and overall read counts is shown for cells prepared as indicated, with cells shown as points shaded based on OAR score calculation. D: temperature for dissociation (warm or cold). P: preservation method (cryopreserved, fixed or fresh).

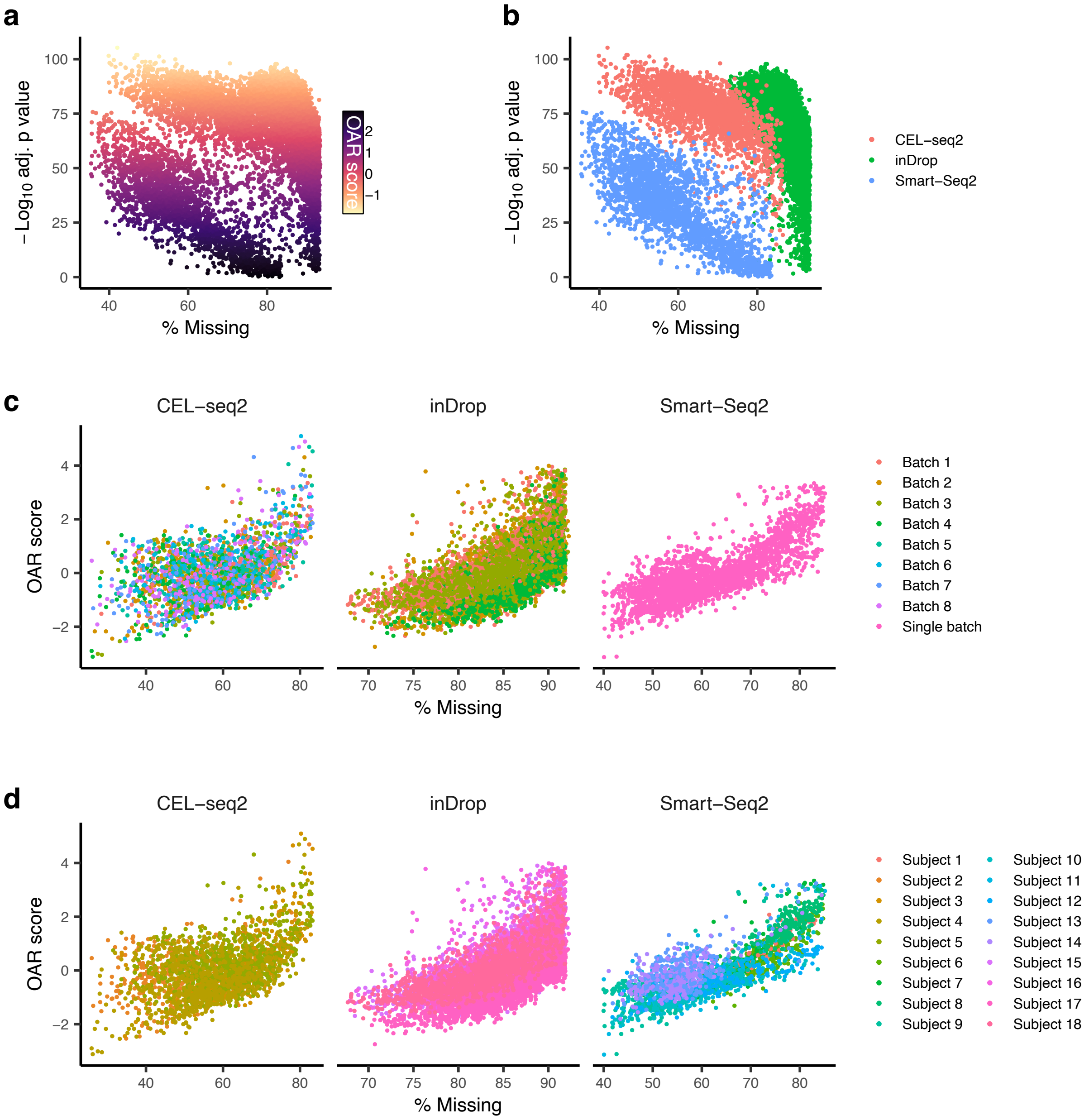
a
b
c
d
−Log10 adj. p value
% Missing
OAR score
CEL−seq2
inDrop
Smart−Seq2
Batch 1
Batch 2
Batch 3
Batch 4
Batch 5
Batch 6
Batch 7
Batch 8
Single batch
Subject 1
Subject 2
Subject 3
Subject 4
Subject 5
Subject 6
Subject 7
Subject 8
Subject 9
Subject 10
Subject 11
Subject 12
Subject 13
Subject 14
Subject 15
Subject 16
Subject 17
Subject 18

***Supplementary Figure 8. OAR scores are robust across protocols and batches.***

**a-d**, OAR scores were calculated on human pancreas cells obtained from three independent studies[5–7] employing separate single cell RNA-sequencing protocols, across several batches of cell preparation and 18 donors. **a,b**, Adjusted Kruskal-Wallis p-values and percent missing genes are shown for all cells when OAR test was applied on all cells simultaneously, with cells as points shaded based on OAR scores (**a**) or for each protocol separately (**b**). **c,d**, OAR scores and percent missing genes in cells when the test was applied to each protocol separately are also shown, with cells as points shaded based on batches (**c**), or donors (**d**).

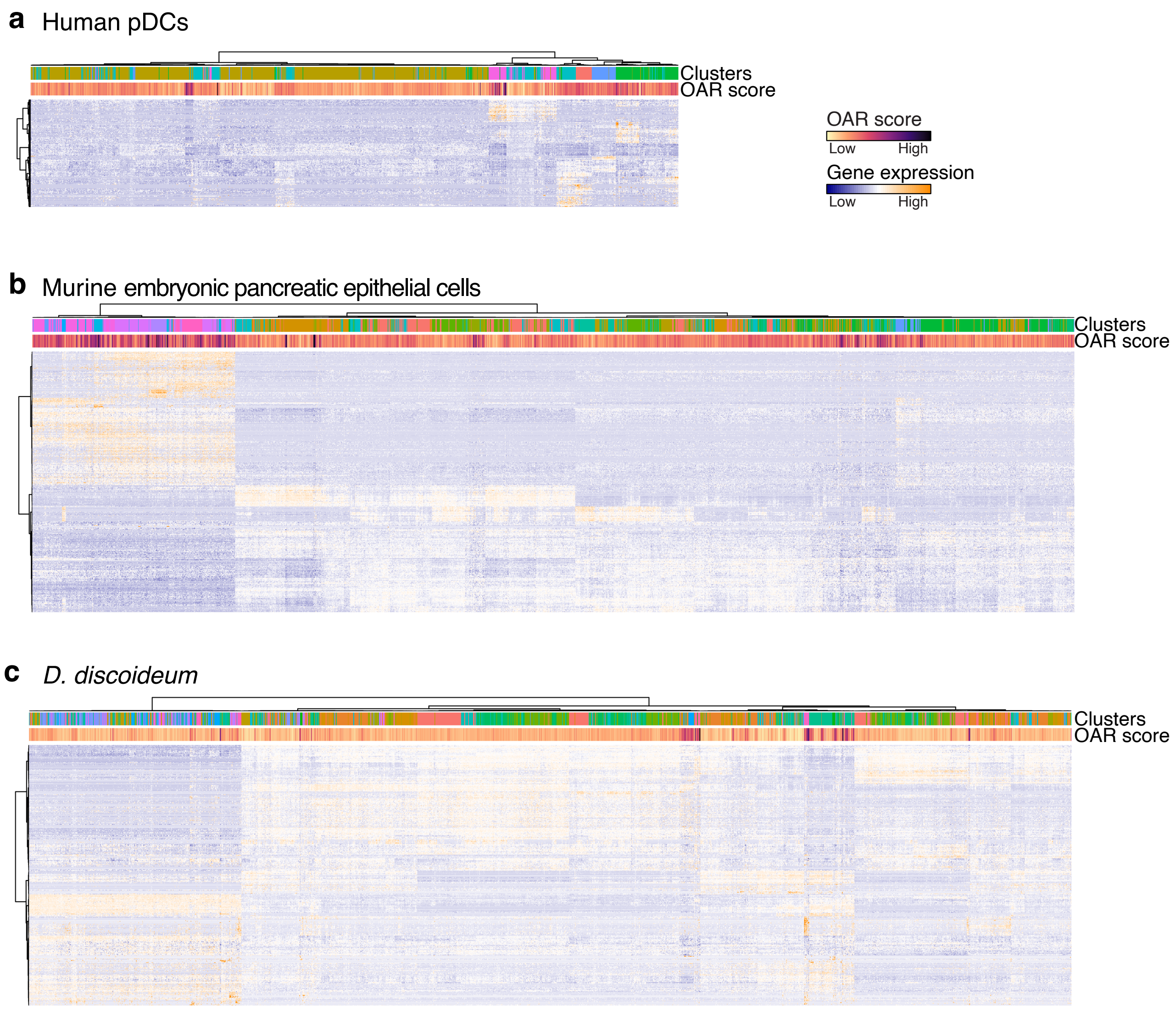

a
Human pDCs
Clusters
OAR score
OAR score
Low
High
Gene expression
Low
High
b
Murine embryonic pancreatic epithelial cells
Clusters
OAR score
c
D. discoideum
Clusters
OAR score

***Supplementary Figure 9. OAR associated genes partially reconstruct cell clustering.***
**a-c**, Row-scaled expression of OAR associated genes (rows) across all cells (columns) are shown, with OAR scores and predicted clusters presented (column annotations), for indicated datasets. Only the top 1000 most significant genes are shown for murine embryonic pancreatic epithelial cells (**b**) and *D. discoideum* (**c**) datasets.

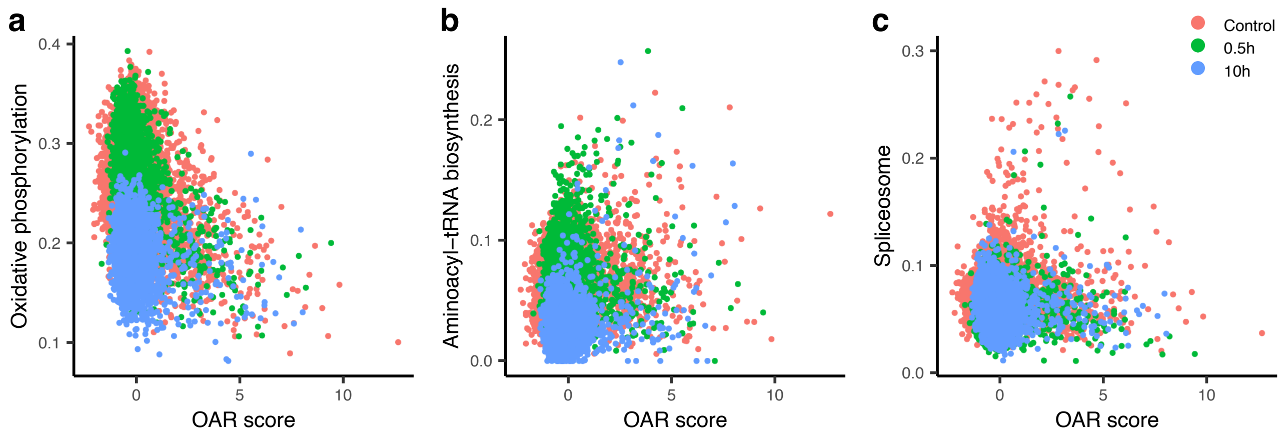
a
Oxidative phosphorylation
0.4
0.3
0.2
0.1
0
5
10
OAR score
b
Aminoacyl−tRNA biosynthesis
0.2
0.1
0.0
0
5
10
OAR score
c
Spliceosome
0.3
0.2
0.1
0.0
0
5
10
OAR score
Control
0.5h
10h

***Supplementary Figure 10. OAR associated genes programs in D. discoideum.***

**a-c**, Indicated gene expression programs enriched in genes associated with OAR score are shown for all cells in *D. discoideum* dataset, with points representing cells colored based on experimental groups.

***Supplementary Video 1. D. discoideum cells decrease mitochondrial activity as they form multicellular aggregates.***

*D. discoideum* cells were starved for 24h then examined for mitochondrial activity with mitochondrial membrane potential ($\Delta\Psi_m$) sensing probe JC-1 via microscopy. Cells forming multicellular aggregates had consistently depolarized mitochondria, evident as JC-1 monomers (510nm excitation / 527nm emission), whereas cells in the periphery displayed high $\Delta\Psi_m$, evident as JC-1 aggregates (585nm excitation / 590nm emission). 3D reconstruction of cellular aggregate representative of 2 independent experiments is shown.

***Supplementary Video 2. Time-lapse of D. discoideum cells decreasing their mitochondrial activity as they form multicellular aggregates.***

*D. discoideum* cells were starved then examined for mitochondrial activity with mitochondrial membrane potential ($\Delta\Psi_m$) sensing probe JC-1 via microscopy, with images collected every 5 minutes between 10-18h of starvation. Cells forming multicellular aggregates had consistently depolarized mitochondria, evident as JC-1 monomers (510nm excitation / 527nm emission). Differential interference contrast (left) and JC-1 monomer fluorescence (right) videos representative of 2 independent experiments are shown.

**Supplementary Note 1 - Comparing OAR scores with heterogeneity and prioritization metrics.**

High OAR scores can identify groups of cells within a dataset that have distinct transcriptional programs that are linked with their biological context. Our method achieves this in a cluster agnostic way that is unique to our knowledge. We examined how OAR scores related to cluster-based metrics of heterogeneity. Cluster labels were assigned based on a resolution chosen to minimize clustering instability. Single cell RNA-sequencing datasets (**Supplementary Table 1**) were used to compare OAR scores with 3 other metrics of cluster heterogeneity:

1. **Silhouette widths** were calculated for each cell using 'silhouette' from cluster v. 2.1.6[23] with the 30 dimensions of the Harmony embedding as input.
2. **Mean square error (MSE)** was selected based on a benchmarking study that found it performed best compared to other distance metrics[24], with a custom function written to calculate this metric. Briefly, we calculate the average expression of each gene across a cell cluster using 'averageColumns' from ezRun[25], then found the MSE between each average gene expression vector, resulting in a matrix of pairwise distances between all clusters.
3. The **number of differentially expressed genes** in a cluster has been proposed as a metric of cluster prioritisation[26]. We used 'FindAllMarkers' from Seurat v.5[27] with default parameters to identify the differentially expressed genes in each cluster, defined as those expressed in at least 50% of the cells in the cluster with an adjusted p-value equal to or less than 0.05, and an average $Log_2$ Fold change equal to or greater than 0.5.

OAR scores highlight transcriptional shifts, which could be associated with cells with a low silhouette width for a specific cluster. We extracted the cluster with the highest median silhouette width from the pDCs (GSE157305), murine embryonic pancreatic epithelial cells (GSE132188), and *D. discoideum* (GSE164010) datasets and examined the relationship between silhouette width and OAR score (**Supplementary Fig. 11**). We found no association between the two, indicating that OAR scores are not indicators of clustering quality.

We then examined the MSE and number of DEGs across 14 macrophage datasets from diverse tissues and biological insults (**Supplementary Fig. 12**). We also calculated OAR scores for each cell and a median OAR score per cluster (**Supplementary Fig. 12**). We found little to no correlation between these metrics of cluster heterogeneity and OAR scores (**Supplementary Fig. 12o**) though it should be noted that median OAR scores do not appropriately capture the outcome of the test we have developed. Notably, OAR scores are not always associated with specific clusters (**Supplementary Fig. 12h-n**).

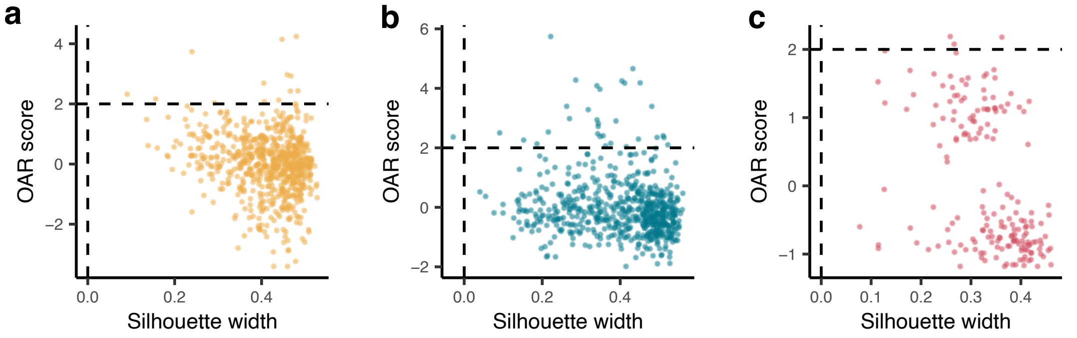


pDCs - Cluster: 0
Median silhouette width: 0.43

Pancreatic epithelial cells - Cluster: 14
Median silhouette width: 0.44

*D. discoideum* - Cluster: 14
Median silhouette width: 0.34

***Supplementary Figure 11. OAR scores calculated in clusters of cells with high silhouette widths.***

**a-c**, Cells from the cluster with the highest median silhouette width were identified within each of the indicated datasets and subsequently an OAR score was calculated for these cells. Points represent cells.

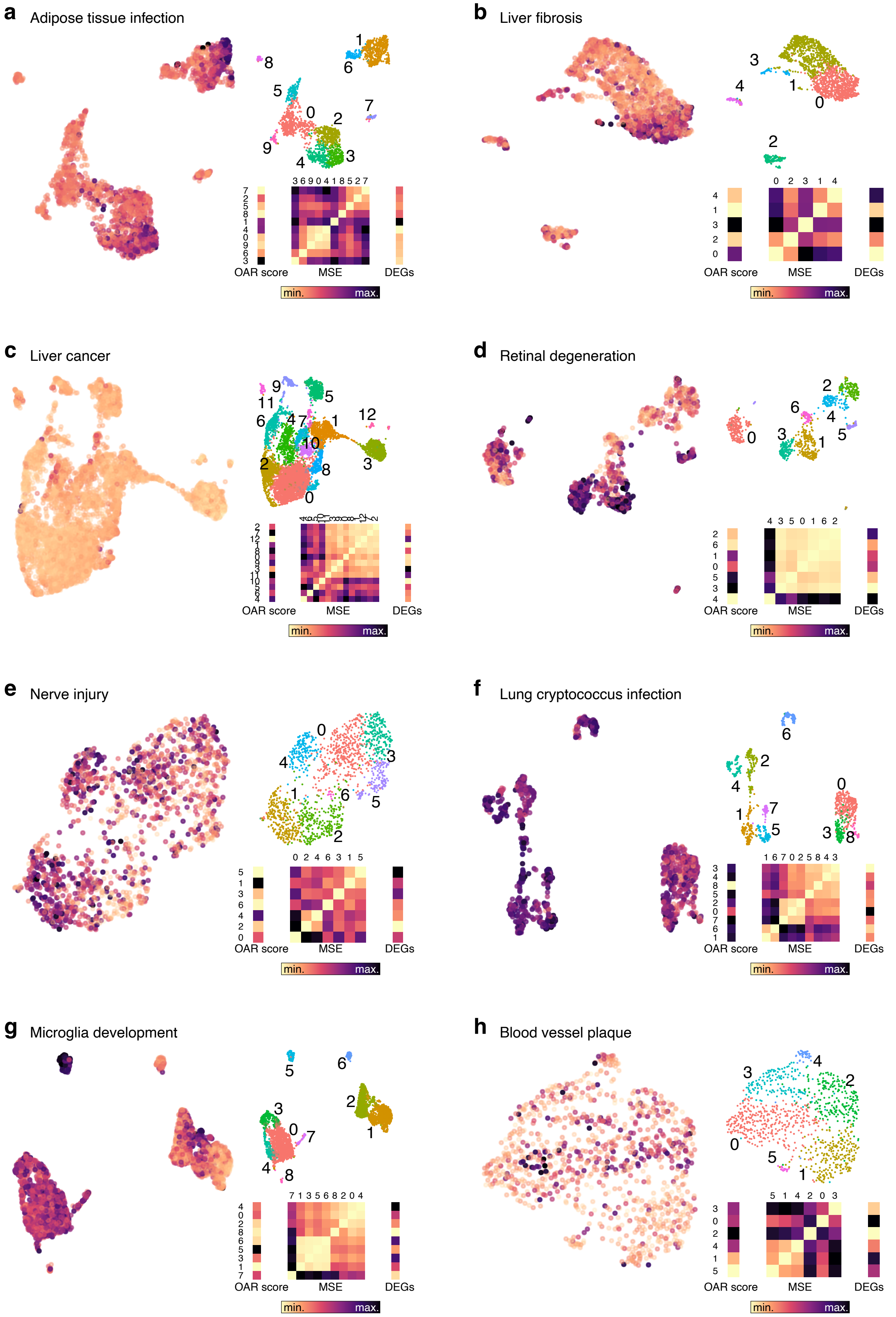
a
Adipose tissue infection
OAR score
MSE
DEGs
min.
max.
b
Liver fibrosis
OAR score
MSE
DEGs
min.
max.
c
Liver cancer
OAR score
MSE
DEGs
min.
max.
d
Retinal degeneration
OAR score
MSE
DEGs
min.
max.
e
Nerve injury
OAR score
MSE
DEGs
min.
max.
f
Lung cryptococcus infection
OAR score
MSE
DEGs
min.
max.
g
Microglia development
OAR score
MSE
DEGs
min.
max.
h
Blood vessel plaque
OAR score
MSE
DEGs
min.
max.

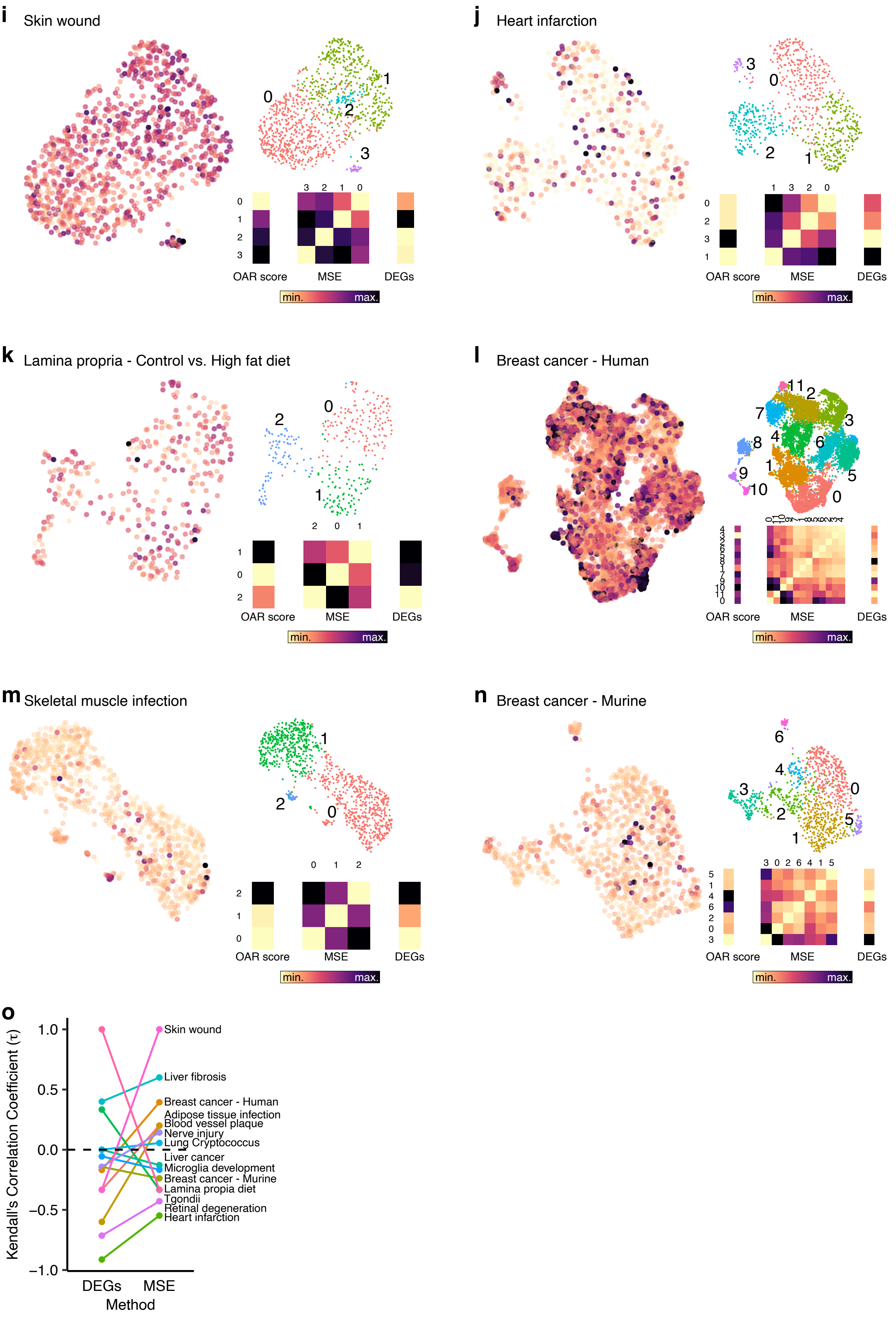

i Skin wound
j Heart infarction
k Lamina propria - Control vs. High fat diet
l Breast cancer - Human
m Skeletal muscle infection
n Breast cancer - Murine
OAR score
MSE
DEGs
min.
max.
o
Kendall's Correlation Coefficient (τ)
1.0
0.5
0.0
−0.5
−1.0
DEGs
MSE
Method
Skin wound
Liver fibrosis
Breast cancer - Human
Adipose tissue infection
Blood vessel plaque
Nerve injury
Lung Cryptococcus
Liver cancer
Microglia development
Breast cancer - Murine
Lamina propria diet
Tgondii
Retinal degeneration
Heart infarction

***Supplementary Figure 12. Comparing OAR scores with cluster based measures of heterogeneity.***

**a-n**, UMAPs showing OAR scores and cluster labels, plus heatmaps showing median OAR scores, mean square error (MSE) and number of differentially expressed genes (DEGs) per cluster for indicated macrophage datasets. **o**, Kendall's correlation coefficient between cluster median OAR scores and DEGs or cumulative MSE for indicated datasets.
